# Molecular and structural mechanism of pan-genotypic HCV NS3/4A protease inhibition by glecaprevir

**DOI:** 10.1101/692392

**Authors:** Jennifer Timm, Klajdi Kosovrasti, Mina Henes, Florian Leidner, Shurong Hou, Akbar Ali, Nese Kurt-Yilmaz, Celia A. Schiffer

## Abstract

Hepatitis C virus (HCV), causative agent of chronic viral hepatitis, infects 71 million people worldwide and is divided into seven genotypes and multiple subtypes with sequence identities between 68 to 82%. While older generation direct-acting antivirals (DAAs) had varying effectiveness against different genotypes, the newest NS3/4A protease inhibitors including glecaprevir (GLE) have pan-genotypic activity. The structural basis for pan-genotypic inhibition and effects of polymorphisms on inhibitor potency were not well known due to lack of crystal structures of GLE-bound NS3/4A or genotypes other than 1. In this study, we determined the crystal structures of NS3/4A from genotypes 1a, 3a, 4a and 5a in complex with GLE. Comparison with the highly similar grazoprevir (GZR) indicated the mechanism of GLE’s drastic improvement in potency. We found that while GLE is highly potent against wild type NS3/4A of all genotypes, specific resistance-associated substitutions (RASs) confer orders of magnitude loss in inhibition. Our crystal structures reveal molecular mechanisms behind pan-genotypic activity of GLE, including potency loss due to RASs at D168. Our structures permit for the first time analysis of changes due to polymorphisms among genotypes, providing insights into design principles that can aid future drug development and potentially can be extended to other proteins.

## Introduction

An estimated 71 million people (∼3.5M in the US) are chronically infected with HCV, which is the leading cause of liver cancer and cirrhosis.^*1*^ There are seven different HCV genotypes (GTs) and multiple subtypes of diverse global distributions with GT1 accounting for ∼50% and GT3 for ∼30% of the global infections.^*2-5*^ Genotypes 1 and 2 have a diverse global distribution; 3 is endemic in South Asia, 4 in the Middle East and Central Africa, 5 in South Africa, 6 in Asia and 7 in central Africa.^*2-5*^ In the last decade the treatment of HCV infection has been revolutionized with direct-acting antivirals (DAAs) including NS3/4A protease inhibitors (PIs),^*6-10*^ but the genetic diversity among genotypes and within a viral population presented a challenge to the development of efficient therapies.

HCV NS3/4A is a bifunctional protein comprised of an N-terminal protease domain and a C-terminal helicase domain. The protease domain (amino acids 1–180) is a serine protease requiring an 11 amino acid peptide from NS4A as a cofactor for folding and activity. The protease is essential for viral maturation, responsible for cleaving the viral polyprotein at various sites (3-4A, 4A4B, 4B5A, and 5A5B). HCV NS3/4A protease sequences vary among the seven genotypes with sequence identities ranging from 68% to 82% (Table S.1). So far, structural and most biochemical studies focused on GT1a, the only GT that allowed structural characterization. Without crystal structures of NS3/4A proteases of the other genotypes, the impact of various polymorphisms and sequence variations, especially those outside the active site, on protease structure, activity or inhibition has not been investigated. Previously, we created a chimeric protease to emulate the inhibition profile of GT3a by substituting three active site polymorphisms (R123T, D168Q and I132L) into GT1a NS3/4A.^*11*^ This GT1a3a chimera largely recapitulated inhibition characteristics of GT3a, and allowed crystal structure characterization. Other than the GT1a3a chimera, no structure of non-GT1a NS3/4A has been determined before and differences among genotypes have been unexplored.

HCV genotypes have varied resistance-associated substitutions (RASs), and susceptibility to DAAs. The 7 FDA approved all-oral DAA combination therapies have varied effectiveness, and especially the earlier combinations can fail against certain genotypes.^*6*^ Fortunately, the three newest oral DAA regimens, Epclusa (sofosbuvir, velpatasvir),^*12*^ Vosevi (sofosbuvir, velpatasvir, voxilaprevir),^*13, 14*^ and Mavyret (pibrentasvir, glecaprevir),^*15, 16*^ are effective against all HCV genotypes with improved sustained virological response (SVR) rates and good tolerance in patients. While Epclusa, which does not contain a PI, is widely used, Mavyret with the latest generation PI glecaprevir (GLE; **Figure 1A**) is the most recommended therapy due to its short 8-week treatment duration and pan-genotypic activity, especially for treatment-naive patients without cirrhosis.^*8-10*^ In clinical studies Mavyret had a cure rate of >98%, and treatment failures of < 1% are primarily reported for patients infected with GT3a.^*17*^ The basis of improved activity of GLE is not readily apparent considering the stark similarity in chemical structure with the earlier PI grazoprevir (GZR), which had lower potency especially against GT3 and certain resistance-associated substitutions (RASs).

**Figure 1:**
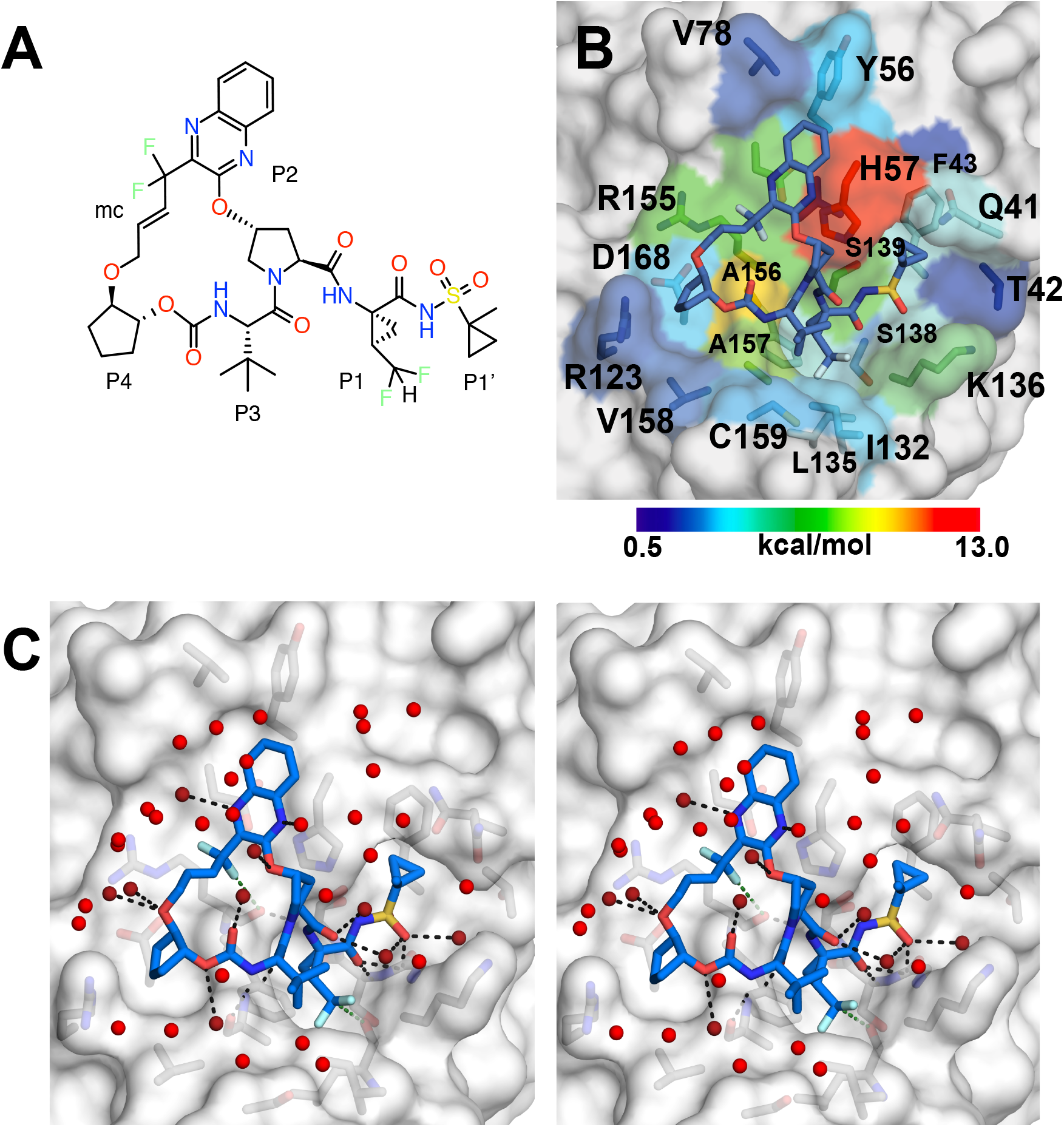
Structure of Glecaprevir. (A) Chemical structure of GLE with the peptide positions labeled. mc = macrocycle (B) GLE bound in the active site of GT1a with the protease surface colored by vdW interactions between GLE and protease; vdW below 0.5 kcal/mol shown in grey. GLE as well as active site side chains are shown as sticks. (C) Stereo image showing the hydrogen bonds formed by GLE and waters coordinated around the ligand. GLE and active side residues are shown in stick representation, waters are shown as red spheres, hydrogen bonds formed by GLE are represented as dashed lines.

While the treatment for HCV infection has significantly improved, one major threat to the clinical efficacy of all currently available anti-HCV drugs is the emergence of drug resistance.^9-10, 18-21^ In fact, single and double RASs can cause resistance to the entire class of protease inhibitors.^*18-20*^ The most common RASs in the NS3/4A protease occur at residues R155, A156, and D/Q168. While substitutions at A156 reduce replicative fitness of the virus drastically, the other RASs do not significantly impair substrate processing and allow robust viral replication. All HCV PIs currently in clinic are peptidomimetic competitive inhibitors and share a common P3–P1’ backbone.^*21*^ Earlier inhibitors showed varying susceptibility to RASs depending on the heterocyclic moiety at the P2 position and the location of the macrocycle.^*22*^ GZR was the first NS3/4A PI with activity against most HCV genotypes and some resistant variants due to its unique binding conformation.^*23, 24*^ Apart from a reduced potency against GT3a, a recent analysis of a clinical study with patients failing GZR monotherapy indicates substitutions at D168 as the main variant responsible for long-term resistance against this compound.^*25*^ D168Q polymorphism is mainly responsible for the reduced PI potency against GT3, and D168E is a common baseline polymorphism in several genotypes with highest prevalence in GT5a.^*17*^ Both GLE and VOX are P2–P4 macrocyclic compounds structurally similar to GZR, and are expected to bind in similar conformations as GZR, supported by their high susceptibility to substitutions at A156.^*26, 27*^ GLE has been reported to lose over 1000-fold potency against RAS A156G and Y56H+Q168R in the GT3a background while Q168R alone conferred 54-fold resistance and Q80R conferred 21-fold resistance against GLE.^*17*^ The Q80K polymorphism, which is outside the active site, is often associated with resistance, primarily in GT1a, GT5a and GT6, and is reported to confer resistance to some PIs and increase the accumulation of other RAS by a mechanism currently not well understood.^*28*^ Thus in addition to the sequence diversity in the active site, polymorphisms among genotypes outside the active site have poorly understood effects on resistance and possibly substrate binding, processing, and protease structure.

Here we report the crystal structures of GLE-bound HCV NS3/4A proteases not only from GT1a but also three other genotypes that had not structurally been characterized. Enzymatic assays revealed GLE is highly potent against wild type NS3/4A protease of all genotypes tested, with inhibition constant in the picomolar range. We also investigated the impact of substitutions at residue 168 on GLE inhibition of the protease. Inhibitor-bound crystal structures of GT1a, 3a, 4a and 5a protease not only reveal the structural basis of pan-genotypic inhibition by GLE, but also structural variation among the genotypes especially outside the active site.

## Methods

### Sequence alignment, construct design and protein expression

Polyprotein sequences of seven HCV genotypes (accession numbers 1a.M62321, 2b.D10988, 3a.D17763, 4a.DQ418788, 5a.AF064490, 6b.D84262, 7b.EF108306) were downloaded and NS3/4A sequences in non-1a genotypes were identified in each polyprotein by alignment with the NS3 and NS4A sequences of genotype 1a used to obtain crystal structures previously (e.g. structure accessible under PDB ID: 5eqq). Alignment and multiple sequence alignments were carried out using BLASTp and COBALT^*29*^ and the multiple alignments were subsequently illustrated using ESPRIPT 3.0.^*30*^ The NS4A peptide sequence of GT2b had to be identified and aligned manually due to low sequence identity. The GT1a construct used for solving structures previously has the 11 amino acid NS4A peptide fused to the N-terminus of NS3 by a 3-amino-acid (SGD) linker.^*31*^ Additionally, it comprised several amino acid substitutions introduced to improve solubility and stability of the protease domain including C159S. GT1a protease with both C159 and S159 were generated. When designing the constructs of the other genotypes, the NS4A peptide was also fused to the N-terminus of NS3 by an SGD-linker and 5 amino acid substitutions were made changing the hydrophobic patch at the surface of the protease domain in helix α1 to hydrophilic residues. Additionally, for genotype 3a the native NS4A sequence was replaced with the NS4A sequence used in the genotype 1a construct. All construct sequences and alignments can be found in the Supplementary Material.

Expression constructs of the non-1a genotypes were ordered from Genescript as codon-optimized sequences for expression in *E. coli*. The GT1a variants D168E, C159S and C159S/D168A as well as the 1a3a chimera were generated previously by site directed mutagenesis starting from the originally optimized GT1a construct.^*11, 23*^ Furthermore, for genotype 4a the SGD linker between NS4A and NS3 was extended to a six-amino acid linker (SGGSGD) using insertion PIPE cloning^*32*^ and confirmed by sequencing. As GT3a protease was more soluble and stable with the C159S mutation, which is close to the S3 and S4 pockets of the active site, we tested the effect of this substitution in the GT1a background where both variants, C159 (GT1a) and S159 (GT1a), could be produced.

All proteases were expressed in *E. coli*, using the same protocol except GT2b. In brief, TB broth supplemented with 30 µg mL^-1^ kanamycin and 0.2% (w/v) D-glucose were inoculated with 25 mL overnight culture (from freshly transformed cells) and grown at 37°C until the OD_600_ reached 0.5–0.6. The temperature was then lowered to 18°C and protein expression was induced with 0.25 mM IPTG for 16 h at 18°C. For GT2b, cells were grown in LB with 30 µg mL^-1^ kanamycin and 0.2 % (w/v) D-glucose and protein expression was induced at OD_600_ 0.5 with 0.2 mM IPTG for 3 h at 37°C. Cells were harvested by centrifugation and stored at −80°C until further processing.

### Protein purification

All proteases and variants were purified using the same protocol and buffer compositions. Briefly, protease-containing *E. coli* cells were thawed, resuspended in Ni-1 buffer (50 mM Phosphate buffer pH 7.5, 500 mM NaCl, 10% (v/v) glycerol, 10 mM imidazole, 2 mM beta-mercaptoethanol) and lysed by passaging twice through a cell disruptor. Cell debris was removed by centrifugation at 10000*xg* for 45 min at 4°C and the supernatant was loaded onto either a 1 mL or 5 mL HisTrap FF crude column (GE Healthcare) equilibrated in Ni-1 buffer using a peristaltic pump. The column was washed in 10 column volumes Ni-1 buffer before connecting to an AKTA Purifier. Protein was eluted with a linear gradient of Ni-2 buffer (50 mM Phosphate buffer pH 7.5, 500 mM NaCl, 10% (v/v) glycerol, 500 mM imidazole, 2 mM beta-mercaptoethanol). Protein-containing fractions were analyzed by SDS-PAGE and protease-containing fractions were pooled, concentrated and loaded onto a Superdex 75 16/60 column (GE Healthcare). The column was equilibrated in either resuspension buffer (50 mM Phosphate buffer pH 7.5, 500 mM NaCl, 10% (v/v) glycerol, 2 mM beta-mercaptoethanol) for subsequent enzymatic activity assays or in crystallization buffer (25 mM MES pH 6.5, 500 mM NaCl, 10% (v/v) glycerol, 2 mM dithiothreitol) for later crystallization experiments. For activity assays, the protein-containing fractions were collected and concentrated to 2 mg mL^-1^ and stored as 20 µL aliquots at −80°C; for crystallization experiments the protein was concentrated to 5–30 mg mL^-1^ and either used fresh, stored at 4°C for a few days or stored in 50 µL aliquots at −80°C.

### Inhibitors

Glecaprevir was purchased from A ChemTek, Inc. (Worcester, MA); the ^1^H- and ^13^-C NMR data of the sample was consistent with the structure. Grazoprevir and danoprevir were synthesized as previously described.^*23*^

### Crystallization

Unless otherwise noted, all protease genotypes and variants were incubated with 3-fold molar excess of inhibitor for 1–3h on ice before setting up the crystallization experiments. Precipitate was removed by centrifugation for 1 min at 10,000*xg* prior to setup. GT1a, GT1a(S159), GT1a(D168E) and GT1a3a chimera in complex with GLE as well as GT1a3a chimera in complex with GZR were screened for crystallization and optimized in hanging drop format with 500 µL well solution per condition and drops containing 1 µL protein–PI solution plus 1 µL well solution. Crystallization was induced by streak seeding with GT1a–danoprevir (DAN) crystal seeds in 1:100 dilution with mother liquor. GT3a–GLE crystals were screened and optimized like the GT1a variants and GT1a3a without the GT1a–DAN seeding, but instead three subsequent rounds of seeding with crystalline precipitate and micro crystals obtained in the GT3a–GLE screen. GT4a and GT5a were initially screened in sitting drop format using the commercially available JCSG*plus* screen and the Crystal Phoenix robot (Art Robbins Instruments) with drops containing 0.3 µL protein plus 0.3 µL well solution. GT4a and GT4a-SGGSGD crystals did not require optimization before data collection, while GT5a crystals were optimized in hanging drop format with 500 µL well solution per condition and drops containing 1 µL protein–PI solution plus 1 µL well solution. Conditions leading to the best diffracting crystals used for structure determination (as well as processing details mentioned below) are summarized in Table S.2.

### Data collection, structure determination and refinement

Datasets for crystals of GT1a and GT1a(D168E) were collected at 100 K without cryo protection; all other crystals were cryo-protected with 10% ethylene glycol in well solution before data were collected. Data of GT1a3a–GZR and GT3a crystals were collected at beamline 23-ID-B at the Advanced Photon Source (Argonne National Laboratory), and autoprocessed at the beamline with gmcaproc. All other datasets were collected on an in-house Rigaku X-ray system with a Saturn 944 CCD detector and processed using HKL3000^*33*^ and their quality was assessed using Xtriage.^*34*^ The GT5a data were further processed with Aimless and the structure was solved using MrBUMP from the CCP4 program suite.^*35*^ The other structures were solved using PHASER^*36*^ with one NS3/4A GT1a chain (PDB ID: 5voj) used as molecular replacement model. All structures were refined using Phenix refine,^*34*^ and manual modeling of protein chains and ligands was carried out in Coot.^*37*^ Final structures were validated using *Molprobity*^*38*^ before deposition to the PDB. Crystallographic data and statistics are summarized in Table S.3.

### Structure Analysis

Structures were viewed, aligned and hydrogen bonding was analyzed using PyMOL (version 2.1).^*39*^ Comparison of inhibitor binding within all GLE-bound structures as well as GT1a in complex with GZR (PDB ID: 3sud)^*23*^ was carried out by aligning the structures on the C_α_ atoms of the active site residues (42-43, 56-58, 81, 132-139, 154-158, 168). 3sud has 4 chains in the asymmetric unit (AU) with minor changes between the chains. While chain C has the lowest B-values, it showed a slightly different conformation of R155 and D168 compared to the other 3 chains. Chain A was chosen for comparison as it displayed the crucial salt bridges formed between R123, D168 and R155. The GT1a3a chimera in complex with GZR had 2 chains per AU and both were used for analysis to account for the lower data quality and resolution of this data set.

For comparison of the structures of different genotypes and GT1a variants, internal C_α_ atom distances were calculated within each structure and distance difference plots of structure pairs were calculated and analyzed as previously described.^*40*^ To evaluate the binding of GZR and GLE to the different genotypes and variants, van-der-Waals interactions were calculated as described previously.^*41*^

### Enzymatic activity and inhibition assays

#### Determination of Michaelis–Menten (K_M_) Constant

K_m_ constants for GT1a and GT1a(D168A) NS3/4A were previously determined.^*22*^ For GT1a(D168E), GT1a3a, GT2b, GT3a, GT4a, GT5a and GT6b the same protocol was followed. Briefly, 20 µM of substrate [Ac-DE-Dap(QXL520)-EE-Abu-γ-[COO]AS-C(5-FAMsp)-NH2] (AnaSpec) was serially diluted into 2x assay buffer [50 mM Tris at pH 7.5, 5% glycerol, 10 mM DTT, 0.6 mM LDAO, and 4% dimethyl sulfoxide]. The assay was initiated by injection of 10 µL protease using a Perkin-Elmer EnVision plate reader, to a final concentration of 20 nM, in a reaction volume of 60 µL. Substrate fluorescence was measured using a Perkin-Elmer EnVision plate reader (excitation at 485 nm, emission at 530 nm). Inner filter effect corrections were applied to the initial velocities, *V*_o_, at each substrate concentration. *V*_o_ versus substrate concentration graphs were globally fit for 3 repeats to the Michaelis–Menten equation to obtain the K_M_ value.

#### Correction for the Inner Filter Effect

The inner filter effect (IFE) for the NS3/4A protease substrate was determined using a previously described method.^*42*^ Briefly, fluorescence end-point readings were taken for substrate concentrations between 0 µM and 20 µM. Subsequently, free 5-FAM fluorophore was added to a final concentration of 25 µM to each substrate concentration and a second round of fluorescence end-point readings was taken. The fluorescence of free 5-FAM was determined by subtracting the first fluorescence end point reading from the second round of readings. IFE corrections were then calculated by dividing the free 5-FAM florescence at each substrate concentration by the free 5-FAM florescence at zero substrate.

#### Enzyme Inhibition Assays

For each assay, 2 nM of GT1a, GT1a(S159), GT1a(D168A), GT1a(D168E), GT2b, GT4a, GT5a, or GT6b and 4 nM of GT1a3a or GT3a NS3/4A protease was pre-incubated at room temperature for 1 h with increasing concentration of inhibitor (DAN, paritaprevir (PTV), GZR or GLE) in 2x assay buffer [50 mM Tris at pH 7.5, 5% glycerol, 10 mM DTT, 0.6 mM Lauryldimethylamine oxide (LDAO), and 4% dimethyl sulfoxide]. Inhibition assays were performed in non-binding surface 96-well black half-area plates (Corning) in a reaction volume of 60 µL. The reaction was initiated by the injection of 5 µL of HCV NS3/4A protease substrate (AnaSpec) using a Perkin-Elmer EnVision plate reader, to a final concentration of 200 nM. The reaction was monitored using a Perkin-Elmer EnVision plate reader (excitation at 485 nm, emission at 530 nm) for 150 reads. Three independent data sets were collected for each inhibitor with each protease construct of interest. The 12 inhibitor concentration points were globally fit to the Morrison equation to obtain the *K*_i_ value using Prism7.

## Results

### Structural characterization of glecaprevir binding to GT1a NS3/4A protease

Glecaprevir (GLE, **Figure** 1A), one of the newest FDA approved HCV NS3/4A protease inhibitors, is a quinoxaline-based P2–P4 macrocyclic compound structurally similar to GZR. We determined the crystal structure of GT1a protease in complex with GLE to 1.73 Å resolution, enabling detailed analysis of the binding mechanism. Similar to the binding mode of GZR, which we previously determined (PDB ID: 3SUD) ^*23*^, GLE spans the S1’–S4 region of the active site with the P2 quinoxaline moiety stacking on the catalytic triad (**Figure** 1B). Compared with GZR, GLE has several modifications at various positions including a 1-methylcyclopropylacylsulfonamide moiety at P1’, a difluoromethyl cyclopropyl amino acid at P1, cyclopentyl at P4, and a modified P2–P4 linker. The P2 quinoxaline is connected to the P4 cyclopentyl moiety *via* a *trans*-2-butenyloxy linker, with difluoro-substitution at the benzylic position. The GLE acylsulfonamide moiety is tightly bound in the oxyanion hole of the S1’ pocket and the P2 quinoxaline is stacked on the catalytic H57 forming cation-π interactions and contacting the catalytic D81. The P4 cyclopentyl, stabilized by the macrocycle, binds in the S4 pocket, formed by R123 and V158. R123 forms the characteristic salt bridge to D168, stabilizing the active site electrostatic network including hydrogen bonding of D168 to N*ε* and N*η* of R155, as observed with other tight-binding ligands.^*23*^

The vdW interactions between GLE and the protease were analyzed (**Figure** 1B) and revealed strong interactions with many residues including A156, R155 and D168, which are sites of reported RAS. Unlike GZR, both GLE and VOX have difluoro substitution at the benzylic position of the macrocycle. GLE interacts with R155 *via* the difluoro group on the macrocyclic linker, with one fluorine making hydrophobic interactions with R155 side chain and the other forming a halogen bond to the main chain oxygen of the same residue. Additionally, GLE forms a hydrogen bond with the protease through the P1 difluoromethyl group. Due to the electronegativity of the two P1 fluorines, the hydrogen of that group is very acidic, seemingly forming strong interactions with the backbone oxygen of residue L135. Additionally, the structure revealed an extensive ordered water network coordinated around the bound GLE at the active site (**Figure** 1C) with hydrogen bonds to water molecules in addition to those formed between GLE and the protease. This is in agreement with the high potency of GLE, and the high degree of order in the crystal.

### Glecaprevir is a potent inhibitor of all NS3/4A genotypes

NS3/4A from six genotypes (1a, 2b, 3a, 4a, 5a and 6b), as well as a GT4a with an extended linker and the 1a3a chimera, were successfully expressed, purified and their enzymatic activity determined. The HCV NS3/4A protease sequences between the genotypes display sequence identities ranging from 68% to 82% (Table S1). Genotypes 1a, 4a, 5a and 6b are closest in sequence to each other (∼81%), with all active site residues conserved, and 7a and 2b are the most divergent to all the other GTs (68-73%). Active site polymorphisms are limited to GT2b with V78A, D79E and I132L; GT3a with R123T, D168Q and I132L; and GT7b with D79S, R123T, I132V and D168Q. The high degree of conservation among all GTs, especially in the active site region, is highlighted when mapped onto the protease structure (**Figure 3A**). All proteases studied were active with K_M_ values for our fluorogenic substrate within one order of magnitude of each other (**Figure S.2**) as expected considering the high sequence conservation among genotypes of the residues comprising the active site.

**Figure 2:**
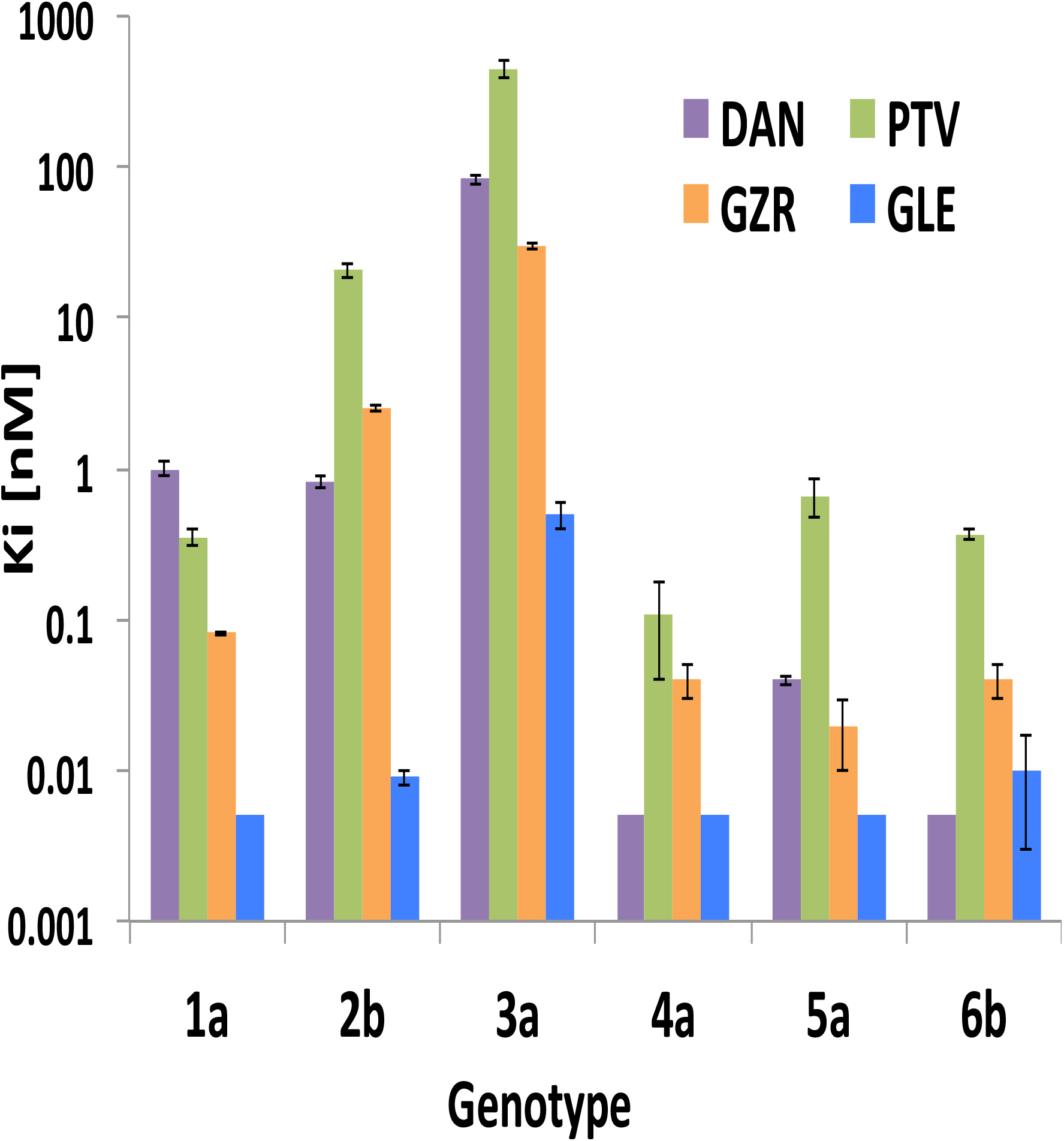
Inhibition of NS3/4A genotypes. Enzyme inhibition constant (K_i_) of danoprevir (DAN, purple), paritaprevir (PTV, green), grazoprevir (GZR, orange) and glecaprevir (GLE, blue) against HCV NS3/4A protease genotypes are plotted in nM on logarithmic scale.

**Figure 3:**
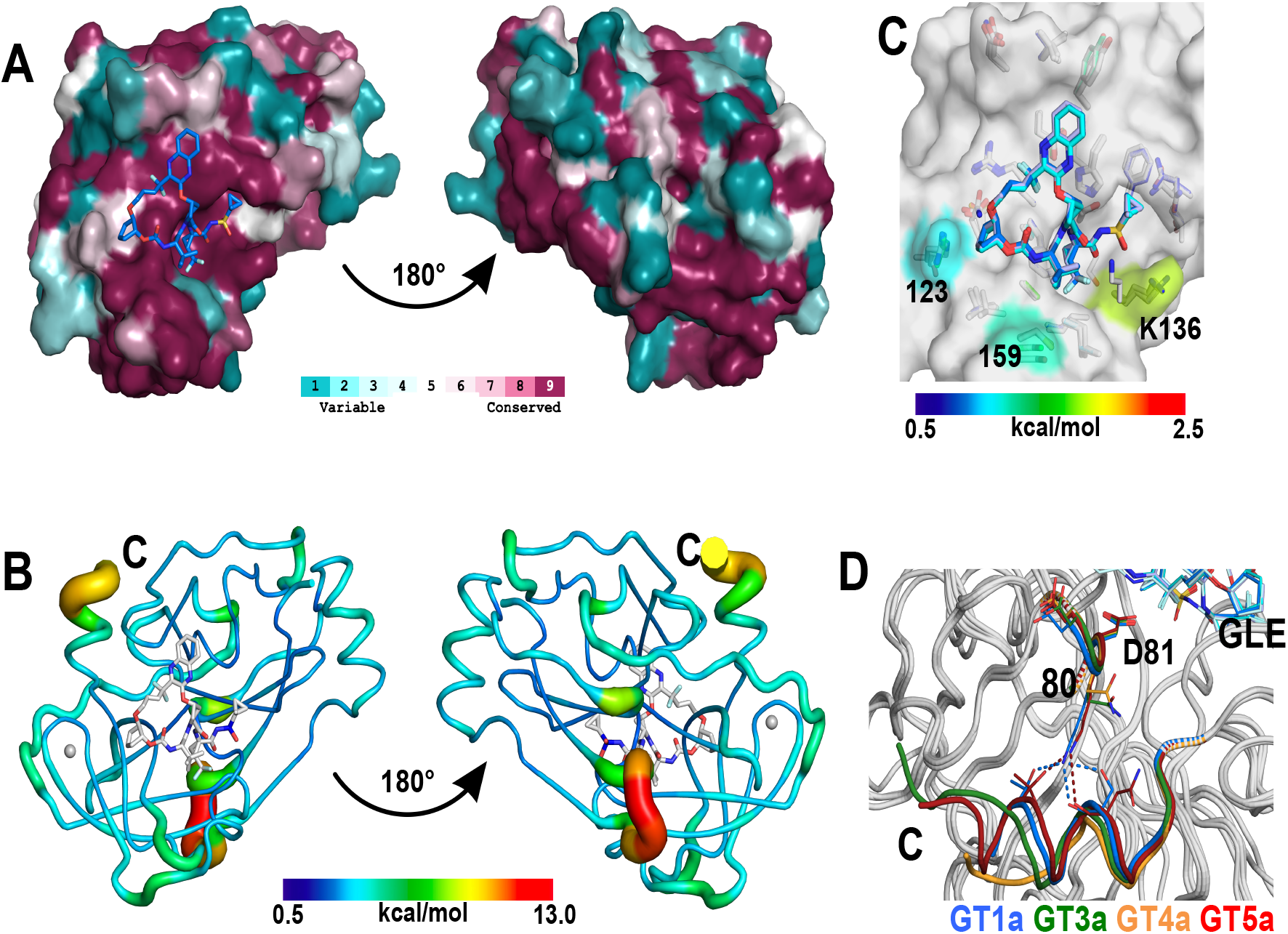
Crystal structures of HCV NS3/4A protease from various genotypes in complex with GLE. (A) GT1a structure with GLE, where the surface is colored by sequence conservation among genotypes. (B) Overall structure of NS3/4A-GLE complex in tube representation with tube thickness and color indicating variation (in intra-molecular Cα distances) over the crystal structures of genotypes 1a, 1a3a, 3a, 4a and 5a. (C) Superposition of the active sites in the crystal structures of 1a, 3a, 4a and 5a protease bound to GLE. The protease surface is colored by differential vdW contacts of GLE to the protease averaged over the genotypes. The surface of residues forming vdW contacts below 0.5 kcal/mol are colored in grey. (D) Overlay of NS3/4A-GLE structures focused on the C-terminus in ribbon representation, with the C-terminal region and loop 78-81 colored by chain with 1a in blue, 3a in green, 4a in yellow and 5a in red. GLE, residues 78-81 as well as residues hydrogen bonded to K80 are shown as sticks with hydrogen bonds shown as dashed lines.

The enzyme inhibition constants (K_i_ values) of PIs including: DAN, PTV, GZR and GLE, against the different genotypes were determined (**Figure 2, Table S.4**). GLE inhibited genotypes 1a, 4a and 5a with a K_i_ value below 5 pM, the detection limit of our assay. Inhibition of GT2b and 6b protease was very efficient with K_i_ of 9 pM and 10 pM, respectively. However, the K_i_ value obtained for GT3a showed a 100-fold decrease in potency (K_i_ = 500 pM) compared to GT1, not surprising considering the similarity of GLE to GZR, which shows a 143-fold decrease in inhibition against GT3a compared to GT1a. This drop in potency was confirmed with the GT1a3a chimera, verifying that the three active site polymorphisms (R123T, D168Q and I132L) are mainly responsible for the less efficient binding of these inhibitors, in agreement with previous findings.^*11*^ While a 100-fold increase in K_i_ for GT3a is significant, GLE still remains highly potent against all NS3/4A genotypes, rendering it a pan-genotypic inhibitor with enzyme inhibition two orders of magnitude better than the previous generation of PIs, and the K_i_ against GT3a comparable to that of GZR against GT1a.

Unlike GLE, the other PIs (DAN, PAR and GZR) have varying inhibition profiles against GTs. While GLE inhibited all GTs except 3a with nearly equal potency, the other three PIs lose several orders of magnitude potency against GT2b (0.82, 20.8 and 2.5 nM for DAN, PTV and GZR respectively). Notably, all PIs inhibited GT 4a, 5a and 6b protease with K_i_ values in the pM range with PTV having the highest K_i_ values. Interestingly, GT1a-S159 was inhibited by GZR slightly weaker than GT1a (K_i_ = 148 pM and 81 pM, respectively). Our constructs for GTs 2b, 4a, 5a and 6b all have the evolutionarily conserved C159 while GT3a NS3/4A has S159, which may somewhat increase the K_i_. Except the active site polymorphisms in GT3a, the reasons for variation in K_i_ values of PIs between the GTs are not straightforward. More long-range effects of non-active site polymorphisms may be responsible for those differences. Nevertheless, GLE performed well against all genotypes, losing potency only against GT3a.

### Overall structure and GLE binding are conserved among genotypes

In addition to the 1a crystal structure, five crystal structures of NS3/4A genotypes in complex with GLE were determined including GT-3a (and 1a3a chimera), two constructs of 4a, and 5a (**Figure** 3). These are the first reported structures of non-1a HCV complexes. Overall, the crystal structures of all NS3/4As are similar and in agreement with the high degree of sequence similarity among genotypes, especially in the secondary structure elements. The active site conformation and GLE binding mode of NS3/4A were conserved in all genotypes. Only subtle changes in vdW interactions of GLE with the proteases could be observed (**Figure 3C and S.7**) as was expected considering the similar K_i_ measured and the high degree of conservation of the active site residues. Residues with the highest divergence in vdW interactions were R/T123, K136 and S/C159. While changes in vdW interactions of K136 with GLE were solely dependent on this residue’s conformation, those of 123 were caused by GT3a-specific R123T polymorphism.

Although the protease structures of the different genotypes are overall very similar, analysis of distance difference plots (**Figure S.5**) revealed that outside the active site, these structures show extensive overall structural plasticity with many regions of the enzyme diverging between 1–1.4 Å (**Figure 3B** and **Figure S.6**) with respect to each other and some loop regions diverging up to 7 Å. Comparison of GT1a and GT1a-S159 structures indicates that C159S substitution caused reduction in vdW interactions with GLE (**Figure**s **S7 and S8**), although the K_i_ remains below the detection limit of our assay. For GZR, a ∼2-fold increase in Ki was measured due to this substitution in GT1a protease, which we expect to have a similar effect in GT3a. Otherwise structural differences were the most pronounced in the linker region between NS4A and NS3, and the C-terminus. 30 residues at the N-terminus of GT4a protease could not be resolved due to disorder in both structures of constructs with two different linker lengths, although the crystals contain the full-length protein (data not shown). As the flexibility is present in both GT4a structures with different linker lengths and fundamentally different crystallization conditions (**Table S.1**), this may be functionally relevant. The same region, although resolved in the GT3a crystals, also shows a higher degree of flexibility compared to the core and active site of the protein indicated by a less well-defined electron density map in the region. Thus, although there is a high level of structural similarity between the genotypes, key regions show altered levels of conformational flexibility.

The C-terminal part of the enzyme shows some differences with potential implications for stability, activity or regulation. The very C-terminus is alpha-helical in GTs 1a and 5a with clear definition of the electron density, while GTs 3a and 4a have a largely disordered C-terminus (**Figure 3D**) with very poor electron density definition. Interestingly, GTs 1a and 5a have lysine at amino acid position 80, coordinating to residues 174 and 178, while GTs 3a and 4a contain a glutamine at position 80, not interacting with the C-terminal region. Coordination by K80 may be stabilizing the C-terminus to a certain extent, with implications for the dynamics between the protease domain and the helicase domain in full-length protein. Previous structures of GT1a with glutamine at position 80 (PDB ID: 3KEE)^*43*^ also display a helical C-terminus, indicating the helix is a feature specific to the genotype and not solely determined by amino acid 80.

### Glecaprevir versus grazoprevir inhibition of NS3/4A protease

GLE inhibits NS3/4A GT1a with a K_i_ at least 42-fold better than GZR. To investigate the reason for such an improvement in potency we compared the structures of GT1a–GLE and GT1a–GZR (PDB ID: 3SUD)^*23*^. Aligned structures immediately reveal differences in PI binding conformation and positioning at the active site, also reflected in analysis of the vdW interactions between protein and inhibitor, illustrated in **Figure** 4A (and **Figure** S.3). In both inhibitors, the quinoxaline moiety stacks on the catalytic residues H57 and D81. However, GLE lacks the methoxy group on the quinoxaline, which in GZR contacts Y56, resulting in reduced vdW contacts with Y56 and ∼0.5 Å shift of the quinoxaline moiety away from V78 and D81 but closer to H57. A *trans* double bond in the GLE macrocycle resulted in a conformational shift of the macrocycle and the P4 cyclopentyl moiety, which better filled the S4 pocket with increased contacts to R123 and D168 compared to GZR. GLE also filled and interacted better with the S1 pocket than GZR due to the difluoromethyl group of the P1 moiety, which forms a particularly strong hydrogen bond with the main-chain carbonyl of L135. Moreover, GLE formed additional electrostatic interactions through the difluoro group in the macrocyclic linker, all remaining direct hydrogen bonds to the protein being identical to those in GZR with the acylsulfonamide moiety tightly bound in the oxyanion hole. The water network around GLE was highly ordered (**Figure** 1C), which was missing entirely in the GZR structure (no waters bound in GT1a–GZR chain A, only one in chain C). While crystallization conditions and crystal characteristics may affect the water network, the generally more ordered environment of GLE in the crystal (within 5 Å of the PI), in addition to improved interactions with the protease, is consistent with the higher potency of GLE compared to GZR.

**Figure 4:**
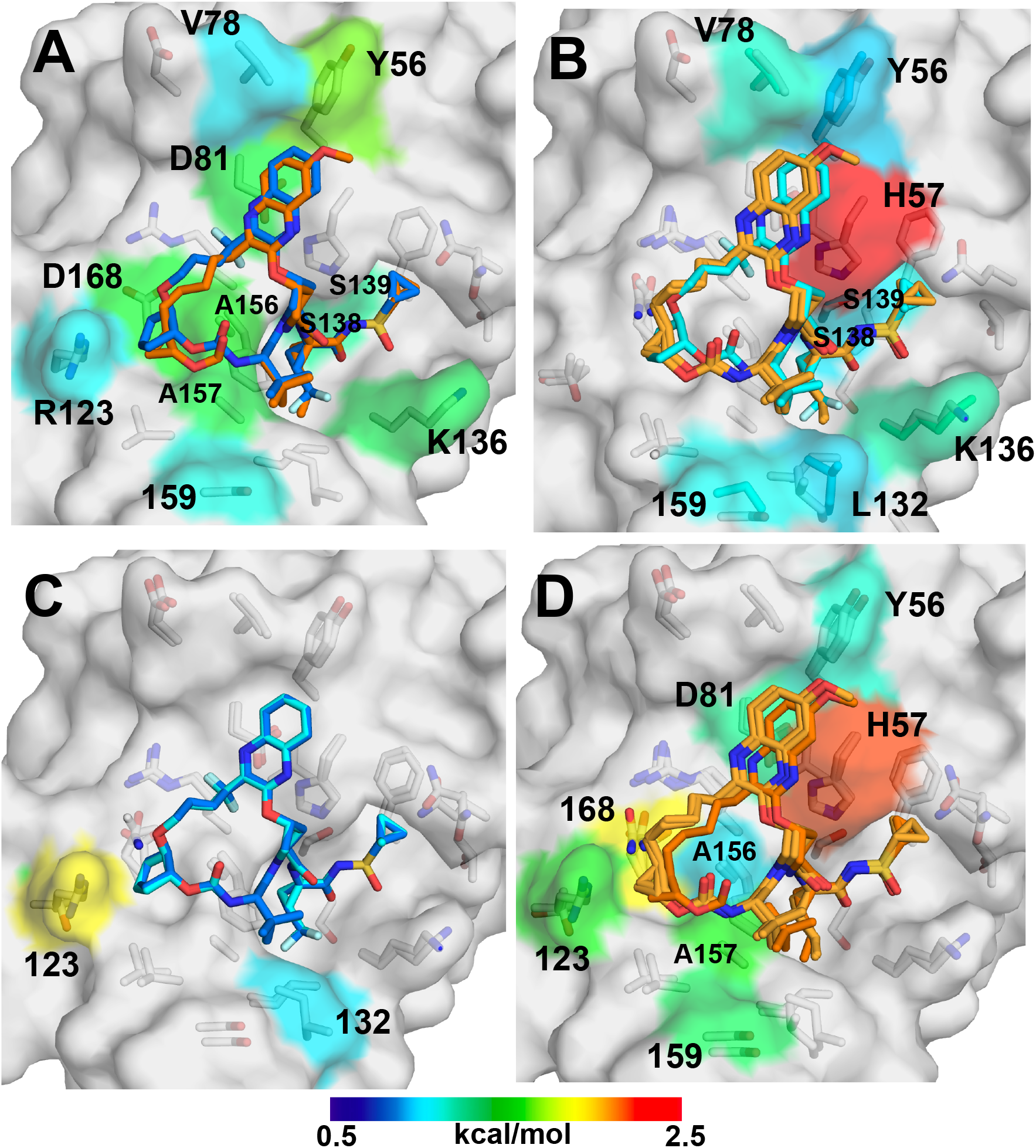
Comparison of the binding modes of GLE and GZR in genotypes 1a and 1a3a. (A) GLE in 1a and 1a3a, (B) GZR in 1a and 1a3a, (C) GLE and GZR in 1a, and (D) GLE and GZR in 1a3a. The inhibitors GLE (blue in 1a, cyan in 1a3a) and GZR (orange in 1a, light orange in 1a3a) as well as the active site residues (grey) are shown as sticks. The surface is colored by differential vdW contacts between the structures compared in each panel.

To address why both GLE and GZR lose substantial potency against GT3a protease, the structures of PI complexes were compared (**Figure** 4B-D). These PIs lost 100-fold and 143-fold potency respectively in the presence of three active site polymorphisms of GT3a, as recapitulated by the 1a3a chimeric construct. Thus, for the detailed comparison the high-resolution co-crystal structure of the 1a3a–GLE complex was used, as GT3a protease failed to crystallize with GZR. Differences in binding and vdW interactions of GZR and GLE were more pronounced in GT1a3a (**Figure** 4B) compared to GT1a. In GT1a3a, GZR had drastically reduced vdW interactions with the catalytic H57 and S139 residues, and pulled out of the S1 pocket, compared to GLE. However, vdW interactions of the protease with the macrocycle and P4 moieties of the two PIs were similar.

Comparing GT1a versus 1a3a NS3/4A (**Figure** 4C), GLE binding conformation was identical except for changes in interactions with residues 123 and 132, which were substituted in 1a3a. Residue 123 is threonine in GT1a3a (GT3a and GT7a) and arginine in all other genotypes. In GT1a, R123 side chain formed a salt bridge to D168, stabilizing the active site conformation, and was positioned in close proximity to GLE. In GT1a3a (and GT3a) structure, the threonine could not form a salt bridge and did not contact GLE, as reflected in much lower vdW interactions. In addition, the D168Q polymorphism in GT1a3a (GT3a) resulted in a loss of hydrogen bond to N*ε* and N*η* of R155 observed for all the other genotypes, as described previously^*11*^, destabilizing the active site conformation. Interestingly, changes in vdW interactions of GLE with the other GT1a3a (GT3a)-specific polymorphisms L132 and Q168 were subtle (less than our 0.5 kcal/mol cutoff) in the crystal structures, but they may affect the dynamic properties of the active site. The high similarity in binding mode and interactions is consistent with sub-nM inhibition by GLE of both GTs.

However, the binding of GZR to the two genotypes is very different (**Figure** 4D). We previously described GZR binding to GT1a NS3/4A in detail (PDB ID: 3SUD)^*23*^. Our new structure of GT1a3a chimera bound to GZR is determined to only 3.5 Å and had two protease chains per asymmetric unit. Both chains have weaker electron density for the GZR macrocycle than the rest of the molecule (**Figure** S.5) and display a 0.5 Å shift of the macrocycle away from the protein compared to GT1a–GZR structure. This may be caused by destabilization due to the Q168 and T123 polymorphisms in GT3a, which fail to form the crucial salt bridge to R155 observed in GT1a. Furthermore, this shift of the macrocycle increased vdW interactions with Q168 and pulled the *tert*-butyl group of GZR further out of S3 pocket, reducing interactions in that region of the active site. The shift in macrocycle further propagated to the quinoxaline moiety, which was shifted 0.5 Å away from H57 reducing the critical cation-π interactions drastically. Hydrogen bonding of GZR with the protein was conserved between GT1a and GT1a3a chimera. Although overall data quality of the 1a3a–GZR structure was not as high as GT1a, taken together the structural differences in GZR binding support and explain the loss of potency against GT3a. Thus structural comparison of GLE and GZR rationalizes the higher potency of GLE against genotypes 1a and 3a.

### Impact of RASs at D168

The effect of substitutions at 168 on inhibition by GLE was characterized for commonly observed RASs. Compared with other genotypes, GT3a Q168 polymorphism is a main contributor to lower potency for most inhibitors including GLE. Additionally, clinical studies showed the polymorphism D168E to be present in patients failing GZR monotherapy.^*25*^ This RAS is a baseline polymorphism commonly found in several genotypes with highest prevalence in GT5a.^*17*^ The highest impact however has been reported for D168A, which is a well-studied RAS conferring resistance to many PIs currently in clinic. GLE loses about 100-fold potency against GT3a and 1a3a chimera, harboring D168Q (**Figure** 5A). However, GLE only loses 2.8-fold potency due to D168E substitution in GT1a, while GZR had a 9.1-fold increase in K_i_. Strikingly, GLE was much more sensitive to the common RAS D168A, showing a 18,000-fold increase in K_i_, compared to 233-fold decrease in potency for GZR. While the increased vdW interactions of GLE with residue 168 in our crystal structures is consistent with susceptibility to D168A, the extent comes as a surprise considering the only 4-fold change in EC_50_ reported using replicon assays.^*26*^

**Figure 5:**
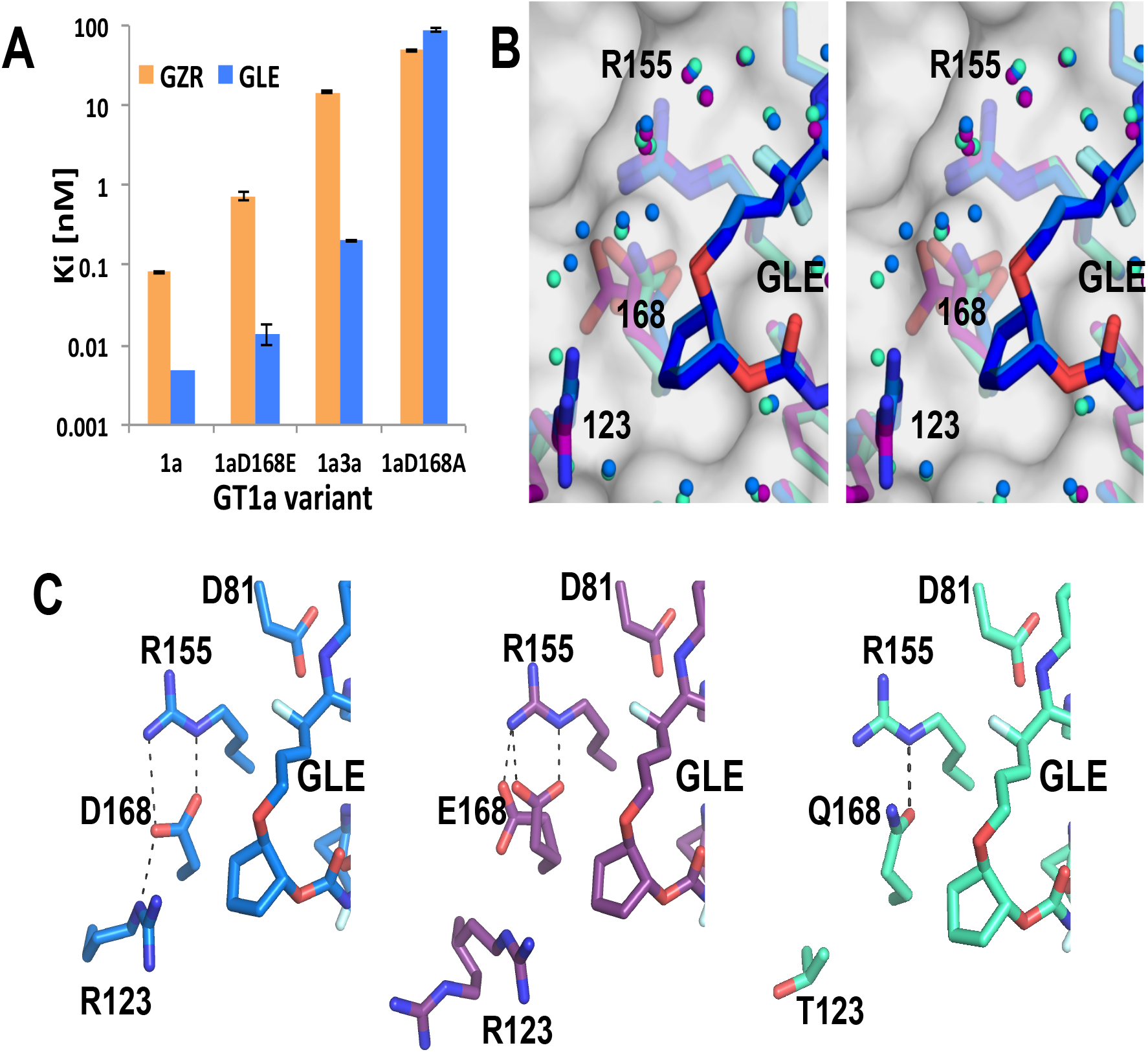
Effect of amino acid 168 on GLE and GZR binding. (A) Inhibition constants (K_i_s) of GZR and GLE against wild-type 1a NS3/4A protease, the clinically relevant 1a-D168E and 1a-D168A variants, and 1a3a chimera (harboring I132L, R123T and D168Q substitutions). (B) Stereo view of the overlaid crystal structures of GT1a (blue), GT1aD168E (purple) and GT1a3a (greencyan) in complex with GLE focused on the region around amino acid 168. Amino acid side chains and GLE displayed as sticks and waters as spheres. (C) Side-by-side view of the GLE-bound structures (GT1a left panel in blue, GT1aD168E middle panel in purple and GT1a3a right panel in greencyan) focused on the region around active site residues 123, 168, 155 and 81 with side chains and GLE in stick representation and hydrogen bonds shown as dashed lines.

To elucidate how GLE tolerates the D168Q and D168E substitutions, crystal structures of GT1a-D168E and GT1a3a chimera in complex with GLE were compared to wild-type GT1a (**Figure** 5B and 5C). The structures overlay very well with no significant changes in GLE binding or the ordered water network. Most of the waters were in identical (or very similar positions) with a few differences in position close to residues 123 and 168 (**Figure** 5B). The structure of GT1a3a–GLE, with T123, shows water molecules occupying the position taken up by R123 in the other two structures. The active site residues have clear electron density in GT1a and 1a3a chimera, revealing differences due to D168Q substitution: In GT1a, D168 forms the characteristic salt bridge to R155 and R123 (**Figure** 5C, left panel) as we previously described.^*23*^ In GT1a3a chimera on the other hand this salt bridge was lost due R123T substitution, and Q168 had a single hydrogen bond to the N*ε* of R155 (**Figure** 5C right panel). These two substitutions and the resulting loss of the salt bridge for stabilization of the active site conformation are likely responsible for the 100-fold loss of affinity of GLE against the 1a3a chimera (and GT3a), which is in agreement with previous studies for other inhibitors.^*11*^

While all active site residues in GT1a and 1a3a chimera are well ordered, the structure of GT1a-D168E in complex with GLE revealed higher flexibility of E168, adopting two alternate conformations (**Figure** 5C middle panel). One rotamer is facing R155 forming hydrogen bonds with N*ε* and N*η*, and the other rotamer was twisted 90° from R155, losing one of the hydrogen bonds. At the same time R123 had less electron density and alternate conformations in line with a high degree of disorder due to lack of coordination with E168. However, while E168 was facing R155, R123 can adopt a conformation very similar to GT1a (D168), potentially forming the characteristic salt bridge between R123, D168 and R155, although in our structures the distances between E168 and R123 were too far for the hydrogen bonds. R155 had the identical conformation observed in GT1a structure and other GLE complexes. Van-der-Waals interactions between GLE and the protein were calculated and compared with GT1a, and the only residue showing changes in vdW above 0.5 kcal/mol was A157. However, these calculations don’t take into account the dynamic nature of the R123 and E168 side chains. Nevertheless, the analysis is in agreement with a smaller loss of GLE potency against GT1a-D168E compared to GT1a3a chimera.

## Discussion

The very potent HCV NS3/4A inhibitor GLE was recently FDA approved as part of a pan-genotypic HCV combination therapy, and in this study we investigated the molecular mechanism of this potency. The structures determined were not only the first co-crystal structures of GLE, but also the first crystal structures of many of the HCV NS3/4A genotypes that had previously remained elusive. These include structures of proteases from GTs 1a, 3a, 4a and 5a, which revealed a high degree of conformational similarity of the active site residues in all genotypes and conservation of intermolecular hydrogen bonding and vdW interactions. With the exception of GT3a, GLE retains potency of 10 pM or better against the NS3/4A proteases of all genotypes. The pan-genotypic activity of GLE potency can be explained by the tight packing against catalytic residues H57 and D81, ordered water structure, and strong electrostatic interactions formed by fluorines in the macrocyle and P1 moiety.

The structural variations between the proteases of different genotypes were limited to regions outside the active site (**Figure 3b**). In the structures the biggest variation concerned the first 30 residues of NS3, which although fairly conserved, are disordered in GT4a and to a lesser extent GT3a. Intriguingly, computational studies suggested the catalytic triad to be less stable in GT3a and even less so in GT4a, compared to GT1a^*44*^ and to be influenced by the cofactor NS4A^*45*^, which is adjacent to the region observed to be disordered in our structures. Another variation among the genotypes involves the loop formed just before catalytic D81 by residues 77–80, NVDQ (**Figure S1**), with substitutions at residue 80 implicated in inhibitor resistance.^*28*^ GT1 and 5 protease sequences frequently have the Q80K polymorphism, while GT2b has an entirely different sequence for residues 77–80, SAEG (**Figure** S1). In our structures the lysine at position 80 had a hydrogen bond to the C-terminal region of the protease (**Figure 3d**) while glutamine 80 does not form this hydrogen bond resulting in a more disordered C-terminal domain. This change may alter the dynamics between protease and helicase domains in addition to potentially influencing the catalytic D81.

Our structures distinguish the PI potency against GT1a from GT3a, with the three active site polymorphisms (R123T, D168Q and I132L). The 100-fold loss in GLE potency against GT3a can be attributed to decreased stability of the active site conformation due to changes in hydrogen bonding pattern and vdW interactions, especially with residue R123T. Although GZR and GLE both lose about 100-150 fold in potency to GT3a compared to 1a, GZR is at least 40 fold weaker against both genotypes, which results in substantial loss in effectiveness against GT3a.

In addition we evaluated RASs D168E and D168A. D168E is a polymorphism in GT5a and an observed RAS associated with GZR^*25*^; and D168A is often observed in patients failing DAA therapy. Both GLE and GZR lost potency against these substitutions, likely due to loss of the ionic network between R123-D168-R155 resulting in decreased vdW interactions and disruption of ordered water network around the inhibitor. Although GLE is a highly potent pan-genotypic inhibitor, drug resistance caused by RASs is still a therapeutic hurdle. Single as well as some double RASs in GT1a and their effect on inhibition have been well studied.^*23, 46-49*^ These include the double mutations Y56H/D168A^*46*^ and the recently observed Y56H/Q168R in GT3a.^*26*^ A156V/T has been shown to cause extremely high levels of resistance to GZR^*22, 23*^ as well as the structurally similar GLE and VOX.^*26, 50, 51*^ RASs at position A156 reduce viral fitness but in combination with other substitutions the fitness is restored.^*18-20, 23, 26, 27, 51, 52*^ Considering the high similarity of all current PIs including GLE, multi-drug resistant variants with RASs at A156 and D168 may become more prevalent in clinic. This threat requires further investigation into molecular mechanisms of resistance and may necessitate development of PIs with altered resistance profiles. Our structural data on GLE binding and non-1a NS3/4A proteases provide crucial insights into potency and resistance, which can be leveraged to devise strategies to design inhibitors with robust pan-genotypic activity.

## Supporting information

Supplementary Matierial

## Acknowledgments

This research was supported by the National Institute of Allergy and Infectious Diseases R01 AI085051. Data was collected at GM/CA@APS which has been funded in whole or in part with Federal funds from the National Cancer Institute (ACB-12002) and the National Institute of General Medical Sciences (AGM-12006). This research used resources of the Advanced Photon Source, a U.S. Department of Energy (DOE) Office of Science User Facility operated for the DOE Office of Science by Argonne National Laboratory under Contract No. DE-AC02-06CH11357. The Eiger 16M detector was funded by an NIH–Office of Research Infrastructure Programs, High-End Instrumentation Grant (1S10OD012289-01A1).

