## Supplementary Matierial for "Molecular and structural mechanism of pan-genotypic HCV NS3/4A protease inhibition by glecaprevir"

### Supplementary Material

#### Protein sequences and alignments

Sequence of GT1a (1a.M62321)
NS4A: GCVVIVGRVVL
NS3:
TAYAQQTRGLLGCIITSLTGRDKNQVEGEVQIVSTAAQTFLATCINGVCWTVYHGAGTRTIASPKGPVIQMYT
NVDQDLVGWPAPQGSRSLTPCTCGSSDLYLVTRHADVIPVRRRGDSRGSLLSPRPISYLKGSSGGPLLCPA
GHAVGIFRAAVCTRGVAKAVDFIPVENLETTMRSP

Sequence of GT2b (2b.D10988)
NS4A: GCISIIGRLHL
NS3:
TAYTQQTRGLLGAIVVSLTGRDKNEQAGQVQVLSSVTQTFLGTSISGVLWTVYHGAGNKTLAGPKGPVTQM
YTSAEGDLVGWPSPPGTKSLDPCTCGAVDLYLVTRNADVIPVRRKDDRRGALLSPRPLSTLKGSSGGPVLC
SRGHAVGLFRAAVCARGVAKSIDFIPVESLDVATRTP

Sequence of GT3a (3a.D17763)
NS4A: GCVVIVGHIEL
NS3:
TAYAQQTRGLLGTIVTSLTGRDKNVVTGEVQVLSTATQTFLGTTVGGVIWTVYHGAGSRTLAGAKHPALQM
YTNVDQDLVGWPAPPGAKSLEPCACGSSDLYLVTRDADVIPARRRGDSTASLLSPRPLACLKGSSGGPVM
CPSGHVAGIFRAAVCTRGVAKSLQFIPVETLSTQARSP

Sequence of GT4a (4a.DQ418788)
NS4A: GSVVIVGRVVL
NS3:
TAYAQQTRGLFSTIVTSLTGRDTNENCGEVQVLSTATQSFLGTAVNGVMWSVYHGAGGKTISGPKGPVNQ
MYTNVDQDLVGWPAPPGVKSLTPCTCGASDLYLVTRHADVVPVRRRGDTRGALLSPRPISTLKGSSGGPLL
CPMGHVAGLFRAAVCTRGVAKAVDFVPVESLETTMRSP

Sequence of GT5a (5a.AF064490)
NS4A: GSVAIVGRIIL
NS3:
TAYAQQTRGVLGAIIVSLTGRDKNEAEGEVQVLSTATQTFLGTCINGVMWTVFHGAGAKTLAGPKGPVVQM
YTNVDKDLVGWPTPPGTRSLTPCTCGSADLYLVTRHADVVPARRRGDTRASLLSPRPISYLKGSSGGPVMC
PSGHVVGVFRAAVCTRGVAKALDFIPVENLETTMRSP

Sequence of GT6b (6b.D84262)
NS4A: GCVVICGRIVT
NS3:
TAYAQQTRGLVGTIVTSLTGRDKNEAEGEVQVVSTATQSFLATTINGVLWTVYHGAGSKNLAGPKGPVCQM
YTNVDQDLVGWPAPLGARSLAPCTCGSSDLYLVTRGADVIPARRRGDTRAALLSPRPISTLKGSSGGPLMC
PSGHVVGLFRAAVCTRGVAKALDFIPVENMDTTMRSP

Sequence of GT7a (7b.EF108306)
NS4A: GSVVVVGRVVL
NS3:
SAYAQQTRGLISTLVVSLTGRDKNETAGEVQVLSTSTQTFLGTNVGGVMWGPYHGAGTRTVAGRGGPVLQ
MYTSVSDDLVGWPAPPGSKSLEPCSCGSADLYLVTRNADVLPLRRKGDGTASLLSPRPVSSLKGSSGGPV
LCPQSHCVGIFRAAVCTRGVAKAVQFVPIEKMQVAQRSP

#### Construct sequences

Sequences of the constructs used are shown below. Hexahistag with thrombin cleavage site and linker to the N-terminus as well as the SGD and SGGSGD linkers are underlined. The amino acid replacements generated for solubility (differing from the database sequences) are highlighted in bold.

Sequence of GT1a construct according to Wittekind *et al.*[1] used for PDB ID 5eqq[2]
MGSSHHHHHHSSGLVPRGSHMASMKKKG**S**VVIVGR**IN**LSGDTAYAQQTRG**EE**GC**QE**TS**Q**TGRDKNQVEG
EVQIVSTA**T**QTFLAT**S**INGV**L**WTVYHGAGTRTIASPKGPV**T**QMYTNVD**K**DLVGW**Q**APQGSRSLTPCTCGSS
DLYLVTRHADVIPVRRRGDSRGSLLSPRPISYLKGSSGGPLLCPAGHAVGIFRAAVCTRGVAKAVDFIPVE**S**L
ETTMRSP

Sequence of GT1a-S159:
MGSSHHHHHHSSGLVPRGSHMASMKKKG**S**VVIVGR**IN**LSGDTAYAQQTRG**EE**GC**QE**TS**Q**TGRDKNQVEG
EVQIVSTA**T**QTFLAT**S**INGV**L**WTVYHGAGTRTIASPKGPV**T**QMYTNVD**K**DLVGW**Q**APQGSRSLTPCTCGSS
DLYLVTRHADVIPVRRRGDSRGSLLSPRPISYLKGSSGGPLLCPAGHAVGIFRAAV**S**TRGVAKAVDFIPVE**S**L
ETTMRSP

Sequence of GT1a-S159;D168A:
MGSSHHHHHHSSGLVPRGSHMASMKKKG**S**VVIVGR**IN**LSGDTAYAQQTRG**EE**GC**QE**TS**Q**TGRDKNQVEG
EVQIVSTA**T**QTFLAT**S**INGV**L**WTVYHGAGTRTIASPKGPV**T**QMYTNVD**K**DLVGW**Q**APQGSRSLTPCTCGSS
DLYLVTRHADVIPVRRRGDSRGSLLSPRPISYLKGSSGGPLLCPAGHAVGIFRAAV**S**TRGVAKAV**A**FIPVE**S**L
ETTMRSP

Sequence of GT1a-D168E:
MGSSHHHHHHSSGLVPRGSHMASMKKKG**S**VVIVGR**IN**LSGDTAYAQQTRG**EE**GC**QE**TS**Q**TGRDKNQVEG
EVQIVSTA**T**QTFLAT**S**INGV**L**WTVYHGAGTRTIASPKGPV**T**QMYTNVD**K**DLVGW**Q**APQGSRSLTPCTCGSS
DLYLVTRHADVIPVRRRGDSRGSLLSPRPISYLKGSSGGPLLCPAGHAVGIFRAAVCTRGVAKAV**E**FIPVE**S**L
ETTMRSP
Sequence of GT1a3a chimera:
MGSSHHHHHHSSGLVPRGSHMASMKKKG**S**VVIVGR**IN**LSGDTAYAQQTRG**EE**GC**QE**TS**Q**TGRDKNQVEG
EVQIVSTA**T**QTFLAT**S**INGV**L**WTVYHGAGTRTIASPKGPV**T**QMYTNVD**K**DLVGW**Q**APQGSRSLTPCTCGSS
DLYLVTRHADVIPVRRRGDSTGSLLSPRPLSYLKGSSGGPLLCPAGHAVGIFRAAV**S**TRGVAKAV**Q**FIPVE**S**L
ETTMRSP

Sequence of GT2b:
MGSSHHHHHHSSGLVPRGSHMASMKKKGCISIIGRLHLSGDTAYTQQTRG**EE**GA**QE**VS**Q**TGRDKNEQAG
QVQVLSSVTQTFLGTSISGVLWTVYHGAGNKTLAGPKGPVTQMYTSAEGDLVGWPSPPGTKSLDPCTCGA
VDLYLVTRNADVIPVRRKDDRRGALLSPRPLSTLKGSSGGPVLCSRGHAVGLFRAAVCARGVAKSIDFIPVE
SLDVA

Sequence of GT3a:
MGSSHHHHHHSSGLVPRGSHMASMKKKG**S**VVIVG**RIN**LSGDTAYAQQTRG**EE**GT**QE**TS**Q**TGRDKNVVTG
EVQVLSTATQTFLGTTVGGVIWTVYHGAGSRTLAGAKHPALQMYTNVDQDLVGWPAPPGAKSLEPCACGS
SDLYLVTRDADVIPARRRGDSTASLLSPRPLAYLKGSAGGPVMCPSGHVAGIFRAAV**S**TRGVAKSLQFIPVE
TLSTQARSP

Sequence of GT4a:
MGSSHHHHHHSSGLVPRGSHMASMKKKGSVVIVGRVVLSGDTAYAQQTRG**EE**ST**QE**TS**Q**TGRDTNENCG
EVQVLSTATQSFLGTAVNGVMWSVYHGAGGKTISGPKGPVNQMYTNVDQDLVGWPAPPGVKSLTPCTCG
ASDLYLVTRHADVVPVRRRGDTRGALLSPRPISTLKGSSGGPLLCPMGHVAGLFRAAVCTRGVAKAVDFVP
VESLETTMRSP

Sequence of GT4a-SGGSGD:
MGSSHHHHHHSSGLVPRGSHMASMKKKGSVVIVGRVVLSGGSGDTAYAQQTRG**EE**ST**QE**TS**Q**TGRDTNE
NCGEVQVLSTATQSFLGTAVNGVMWSVYHGAGGKTISGPKGPVNQMYTNVDQDLVGWPAPPGVKSLTPC
TCGASDLYLVTRHADVVPVRRRGDTRGALLSPRPISTLKGSSGGPLLCPMGHVAGLFRAAVCTRGVAKAVD
FVPVESLETTMRSP

Sequence of GT5a:
MGSSHHHHHHSSGLVPRGSHMASMKKKGSVAIVGRIILSGDTAYAQQTRG**EE**GA**QE**VS**Q**TGRDKNEAEGE
VQVLSTATQTFLGTCINGVMWTVFHGAGAKTLAGPKGPVVQMYTNVDKDLVGWPTPPGTRSLTPCTCGSA
DLYLVTRHADVVPARRRGDTRASLLSPRPISYLKGSSGGPVMCPSGHVVGVFRAAVCTRGVAKALDFIPVE
NLETTMRSP

Sequence of GT6b:
MGSSHHHHHHSSGLVPRGSHMASMKKKGCVVICGRIVTSGDTAYAQQTRG**EE**GT**QE**TS**Q**TGRDKNEAEG
EVQVVSTATQSFLATTINGVLWTVYHGAGSKNLAGPKGPVCQMYTNVDQDLVGWPAPLGARSLAPCTCGS
SDLYLVTRGADVIPARRRGDTRAALLSPRPISTLKGSSGGPLMCPSGHVVGLFRAAVCTRGVAKALDFIPVE
NMDTTMRSP

#### Supplementary tables

**Table S.1: Sequence identity of NS3/4A protease domains on amino acid level calculated in %.** Sequence identity was determined after multiple alignment using Constraint-based Multiple Alignment Tool (Cobalt) https://www.ncbi.nlm.nih.gov/pubmed/17332019.

| | **1a** | **2b** | **3a** | **4a** | **5a** | **6b** | **7a** |
|---|---|---|---|---|---|---|---|
| **HCV.1a.M62321** | - | 69 | 76 | 81 | 82 | 82 | 69 |
| **HCV.2b.D10988** | 69 | - | 68 | 71 | 72 | 71 | 69 |
| **HCV.3a.D17763** | 76 | 68 | - | 74 | 77 | 78 | 73 |
| **HCV.4a.DQ418788** | 81 | 71 | 74 | - | 79 | 81 | 72 |
| **HCV.5a.AF064490** | 82 | 72 | 77 | 79 | - | 83 | 70 |
| **HCV.6b.D84262** | 82 | 71 | 78 | 81 | 83 | - | 68 |
| **HCV.7b.EF108306** | 69 | 69 | 73 | 72 | 70 | 68 | - |

**Table S.2: Crystallization conditions leading the best diffracting crystals used for structure determination.** Cryo-protection and information about processing is added to the table.

| **Protein HCV NS3/4A** | **Ligand** | **Crystal condition and Temperature** | **Cryo** | **Collection and processing** |
|---|---|---|---|---|
| GT1a S159<br>20 mg/mL | Glecaprevir<br>3 mM | 0.1 M MES pH 6.5<br>2% $(NH_4)_2SO_4$<br>20% PEG 3350<br>Seeding with GT1a-danoprevir<br>RT | 10% EG | In house,<br>03.26.18<br>HKL3000, Phenix:<br>Xtriage, Phaser MR,<br>Phenix refine |
| GT1a C159<br>17.5 mg/mL | Glecaprevir<br>3 mM | 0.1 M MES pH 6.5<br>5% $(NH_4)_2SO_4$<br>22% PEG 3350<br>2 μM $ZnCl_2$<br>1 mM TCEP<br>Seeding with GT1a-danoprevir<br>RT | none | In house,<br>12.05.18<br>HKL3000, Phenix:<br>Xtriage, Phaser MR,<br>Phenix refine |
| GT1a C159;D168E<br>17.5 mg/mL | Glecaprevir<br>3 mM | 0.1 M MES pH 6.5<br>2% $(NH_4)_2SO_4$<br>23% PEG 3350<br>2 μM $ZnCl_2$<br>1 mM TCEP<br>Seeding with GT1a-danoprevir<br>RT | none | In house,<br>11.28.18<br>HKL3000, Phenix:<br>Xtriage, Phaser MR,<br>Phenix refine |
| GT1-3 chimera<br>21.6 mg/mL | Grazoprevir<br>4 mM | 0.1 M MES pH 6.5<br>5 % $(NH_4)_2SO_4$<br>24% PEG 3350<br>Seeding with GT1a-danoprevir<br>RT | 10% EG | APS beamline 23 ID-D<br>06.24.18<br>Autoprocessed with<br>gmcaproc, Phenix:<br>Xtriage, Phaser MR,<br>Phenix refine |
| GT1-3 chimera<br>22.3 mg/mL | Glecaprevir<br>3 mM | 0.1 M MES pH 6.5<br>1% $(NH_4)_2SO_4$<br>22% PEG 3350<br>Seeding with GT1a-danoprevir<br>18 C | 10% EG | In house,<br>04.04.18<br>HKL3000, Phenix:<br>Xtriage, Phaser MR,<br>Phenix refine |
| GT3a (S159)<br>7 mg/mL | Glecaprevir<br>3 mM | 0.1 M MES pH 6.5<br>5% $(NH_4)_2SO_4$<br>10 mM $Mg_2SO_4$<br>16% PEG 3350<br>Three rounds of seeding with<br>GT3a-glecaprevir<br>RT | 10% EG | APS beamline 23 ID-D<br>06.24.18<br>Autoprocessed with<br>gmcaproc, Phenix:<br>Xtriage, Phaser MR,<br>Phenix refine |
| GT4a<br>11 mg/mL | Glecaprevir<br>3 mM | 10% PEG 1000<br>10% PEG 8000<br>** JCSG Plus<br>RT | 10% EG | In house,<br>06.11.18<br>HKL3000, Phenix:<br>Xtriage, Phaser MR,<br>Phenix refine |
| GT4a-SGGSGD<br>21 mg/mL | Glecaprevir<br>3 mM | 0.2 M NaCl<br>0.2M HEPES pH 7.0<br>20% PEG 6000<br>** PACT Premier<br>RT | 10% EG | In house, 08.10.18<br>HKL3000, Phenix:<br>Xtriage, Phaser MR,<br>Phenix refine |
| GT5a<br>18.8 mg/mL | Glecaprevir<br>3 mM | 0.2 M TMAO<br>0.1 M Tris pH 8.5<br>16% PEG 2000 MME<br>RT | 10% EG | In house,<br>07.03.18,<br>HKL3000, CCP4:<br>Aimless, MrBUMP,<br>Phenix: Phenix refine |

** Directly from 96 well screen without further optimization

**Table S.3: Crystallographic data and statistics.** The majority of the values in the table were generated using Phenix 'generate Table 1' tool, the remaining values were filled in manually. *Table located in a separate Excel file.*

**Table S.4:** $K_i$ values of danoprevir (DAN), paritaprevir (PTV), grazoprevir (GZR) and glecaprevir (GLE) for the different genotypes and GT1a variants.

| | **$K_i$ (DAN) [nM]** | **$K_i$ (PTV) [nM]** | **$K_i$ (GZR) [nM]** | **$K_i$ (GLE) [nM]** |
|---|---|---|---|---|
| **GT1a** | | | 0.081 ±0.002 | < 0.005 |
| **GT1a-S159** | 1 ± 0.1 | 0.35 ± 0.04 | 0.21 ± 0.03 | < 0.005 |
| **GT1a-S159;D168A** | 199 ± 64 | 297 ± 23 | 49 ± 1.6 | 90 ± 6 |
| **GT1a-D168E** | n.d. | n.d. | 0.74 ± 0.09 | 0.014 ± 0.004 |
| **GT1a3a** | 47.9 ± 1.9 | 63.5 ± 21.8 | 14.8 ± 0.6 | 0.2 ± 0.002 |
| **GT2b** | 0.82 ± 0.08 | 20.8 ± 2.3 | 2.5 ± 0.1 | 0.009 ± 0.001 |
| **GT3a** | 82.8 ± 6.8 | 453 ± 62 | 30 ± 1.9 | 0.5 ± 0.01 |
| **GT4a** | < 0.005 | 0.11 ± 0.07 | 0.04 ± 0.01 | < 0.005 |
| **GT5a** | 0.04 ± 0.003 | 0.66 ± 0.19 | 0.02 ± 0.01 | < 0.005 |
| **GT6a** | < 0.005 | 0.37 ± 0.03 | 0.04 ± 0.01 | 0.01 ± 0.007 |

#### Supplementary Figures

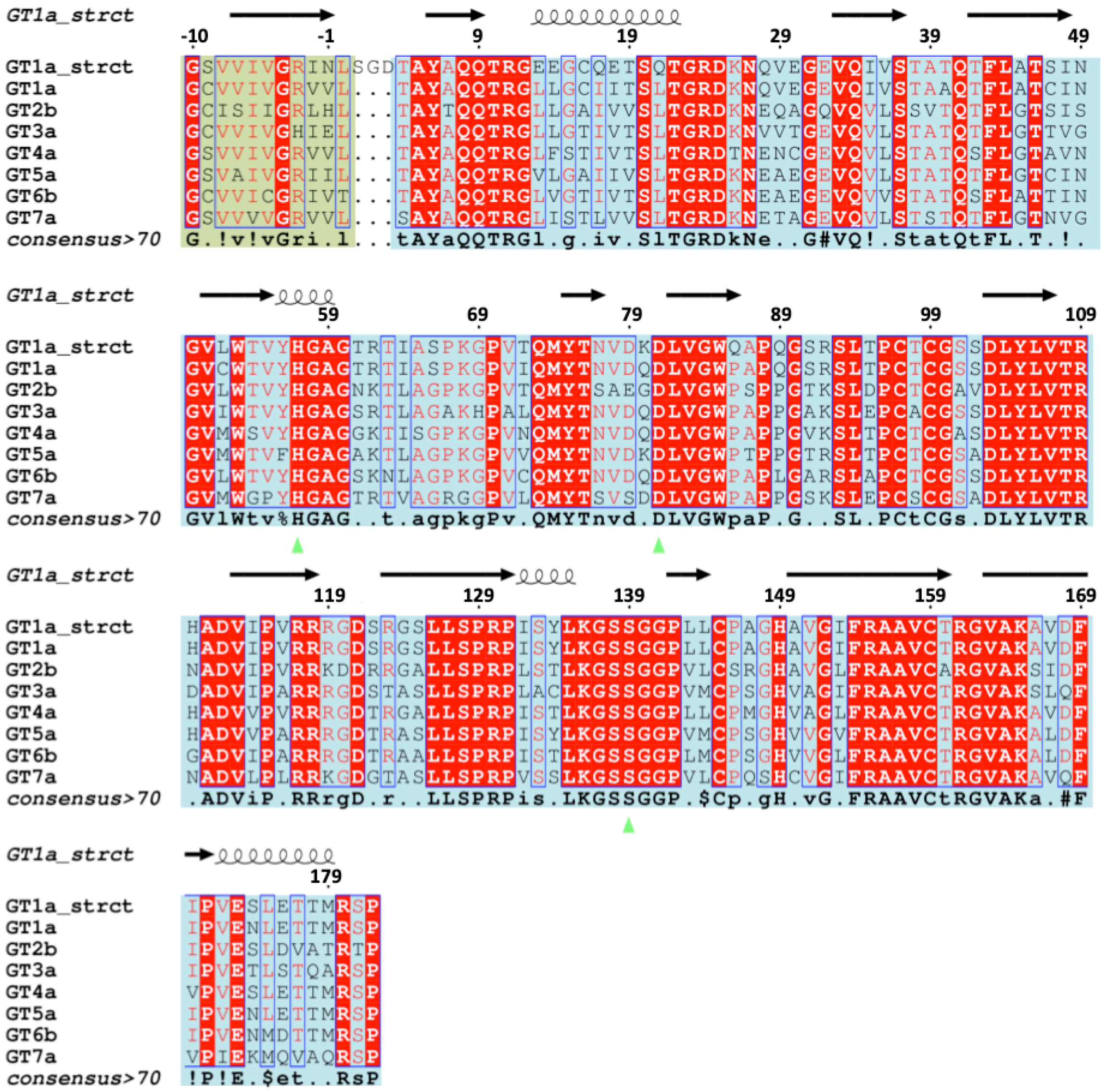


**Figure S.1: Alignment of NS3/4A genotypes.** Sequences (our expression construct with the NS4A peptide fused to the N-terminus of NS3 protease domain *via* a three-amino acid (SGD) linker, genotype sequences of accession numbers 1a.M62321, 2b.D10988, 3a.D17763, 4a.DQ418788, 5a.AF064490, 6b.D84262, 7b.EF108306) were aligned by BLASTp and Constraint-based Multiple Alignment Tool (Cobalt) https://www.ncbi.nlm.nih.gov/pubmed/17332019, and the multiple alignments were subsequently illustrated using ESPRIPT 3.0. NS3 has a pale blue background; NS4A was marked with a pale green background. Amino acids conserved in all sequences are emphasized with red; the green arrows highlight the residues of the catalytic triad.

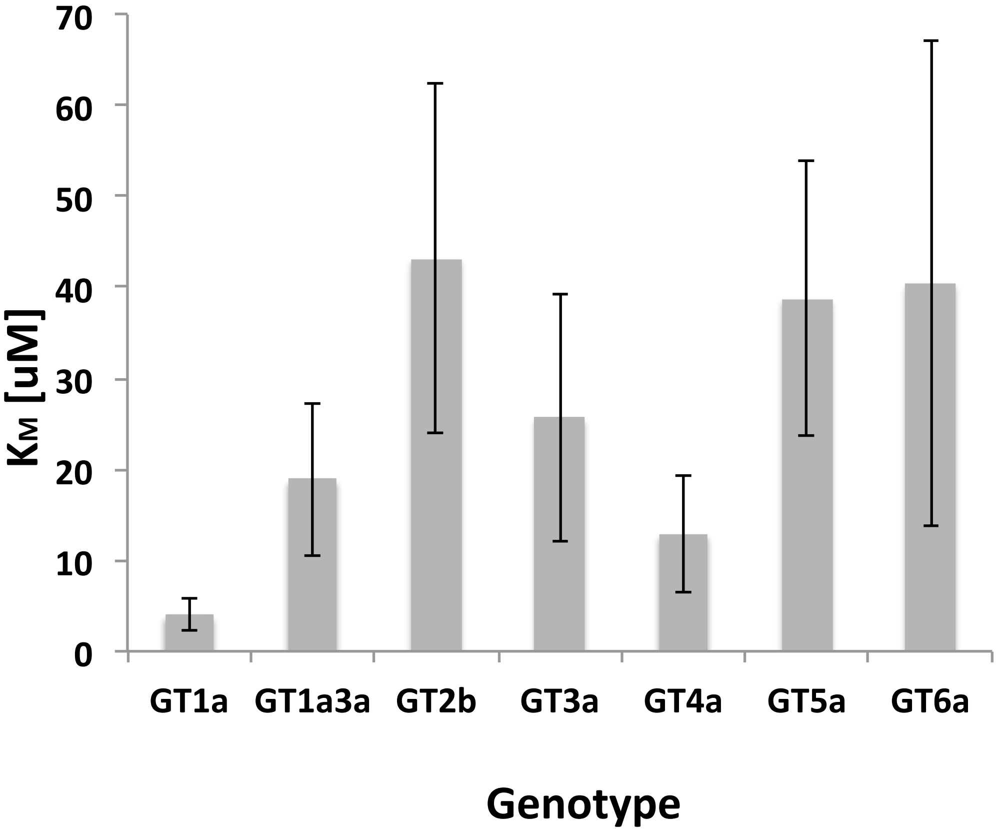


**Figure S.2: Activity of the different genotypes**. $K_M$ data for the different genotypes, including GT1a3a chimera, are shown in µM.

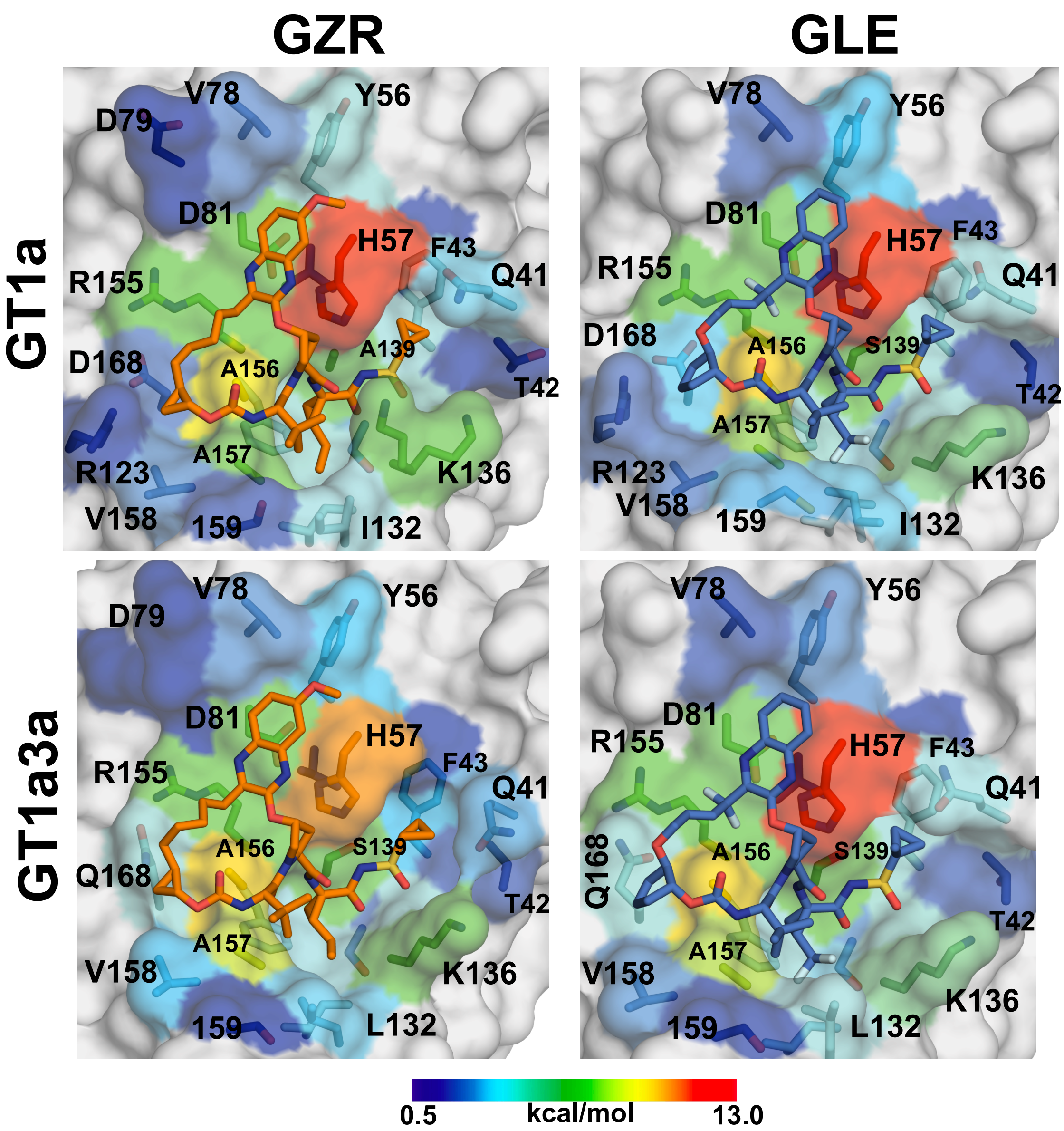


**Figure S.3: Van-der-Waals interactions between NS3/4A and PIs.** Structures of GT1a, non-1a GTs and GT1a3a chimera and GT1aD168E in complex with GLE and GZR were aligned and displayed in Pymol. The Lennard Jones potentials (vdW) were displayed as color gradient on the calculated surface of GT1a and/or GT1a3a. All residues with vdW below 0.5 kcal/mol are shown in grey.

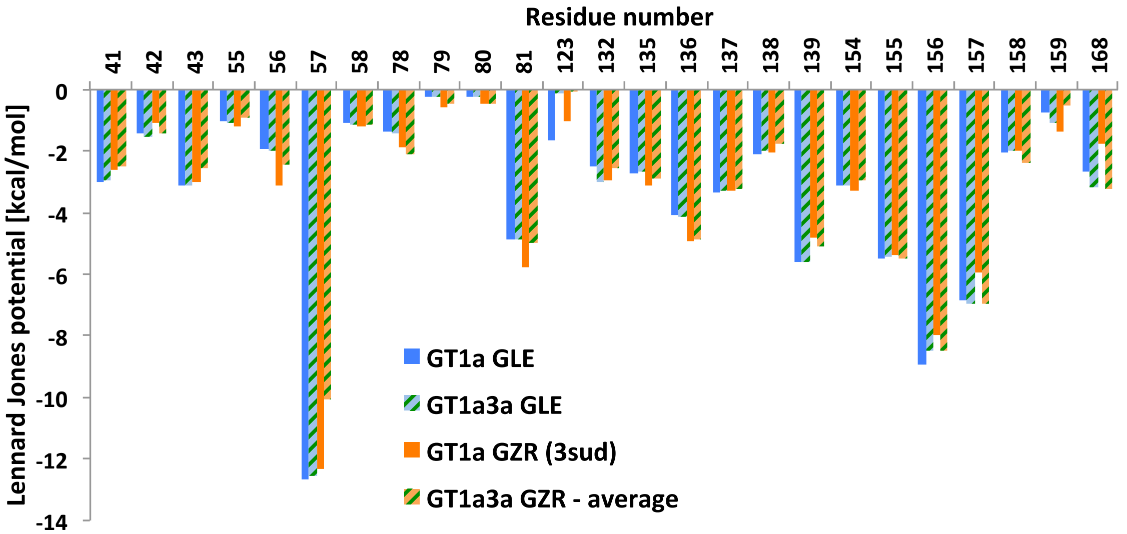


**Figure S.4: Van-der-Waals interactions between NS3/4A and PIs.** The vdW interactions between protease and PIs were calculated per residue and values above 0.5 kcal/mol were plotted. For GT1a_GZR chain A was used for the analysis, while for GT1a3a_GZR both chains were used to calculate an average to account for the lower data quality and resulting potential for error.

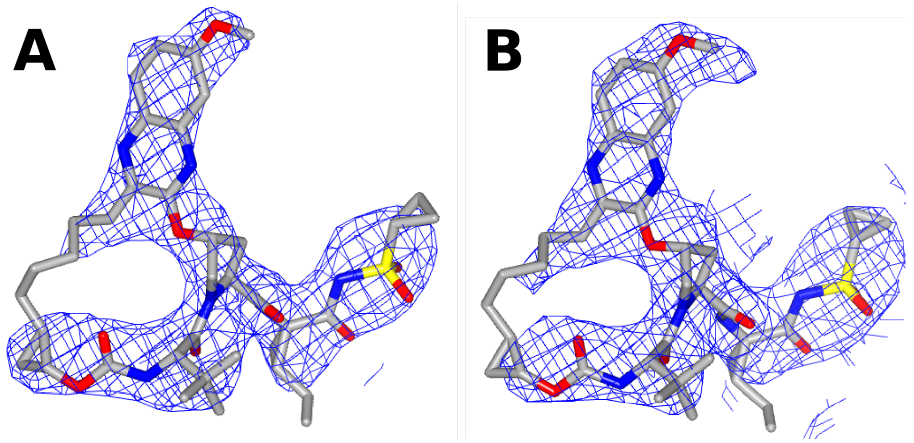


**Figure S.5: Electron density of GZR in GT1a3a chimera.** Shown are the 2Fo-Fc maps contoured at 1 σ of GZR in chain A (A) and chain B (B), clipped around GZR. This figure was made using CCP4*mg*.[3]

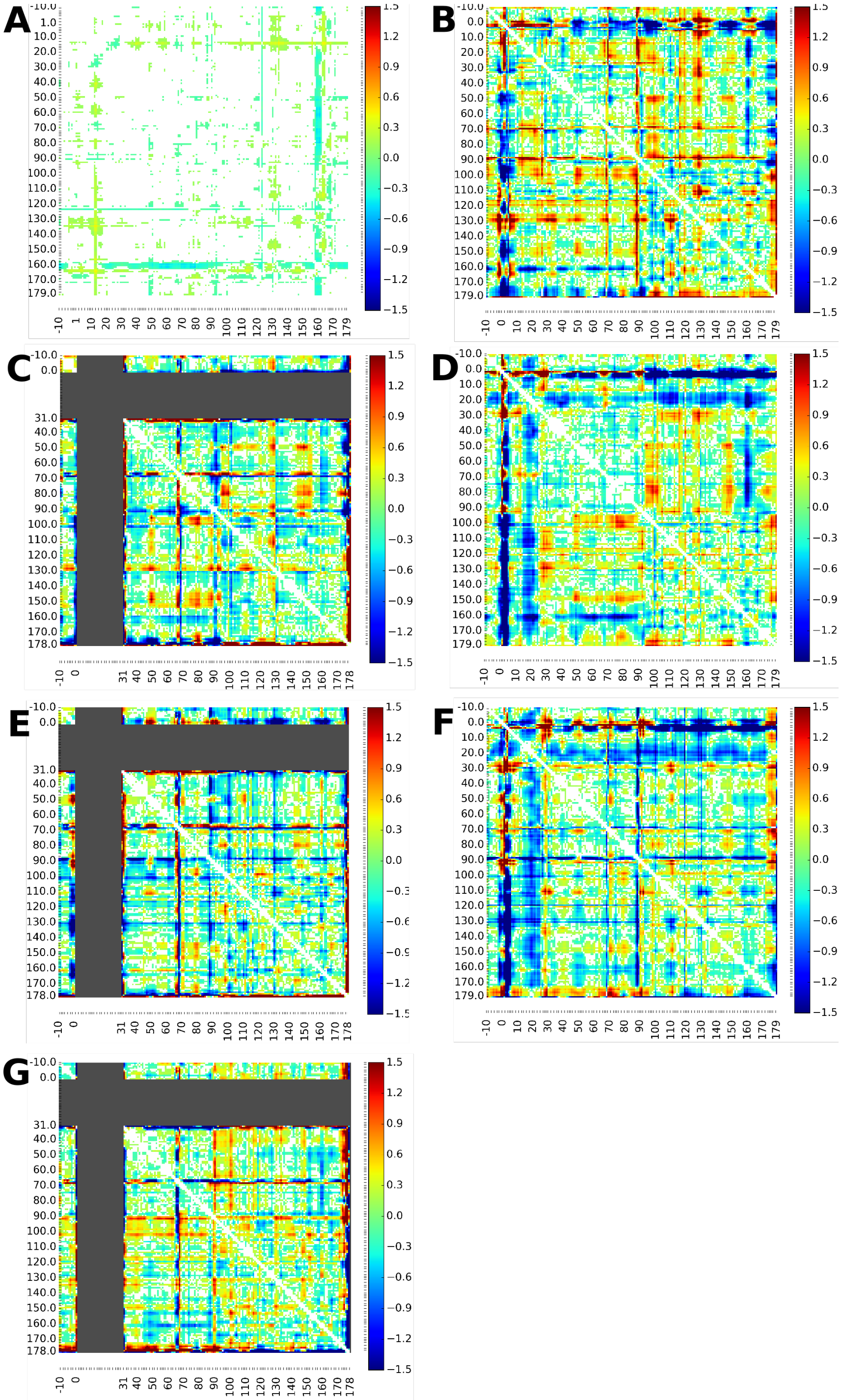


**Figure S.6: Distance difference plots of NS3/4A structures in complex with GLE.** The intra-atom distances of Cα atoms in each structure were calculated and compared pair-wise. The plots represent changes in coordinates per residue. In the GT4a structure, residues 1-30 are missing and hence could not be compared; those non-compared residues are represented in grey. (A) GT1a compared to GT1a3a chimera; (B) GT1a compared to GT3a; (C) GT1a compared to GT4a; (D) GT1a compared to GT5a; (E) GT3a compared to GT4a; (F) GT3a compared to GT5a; and (G) GT4a compared to GT5a.

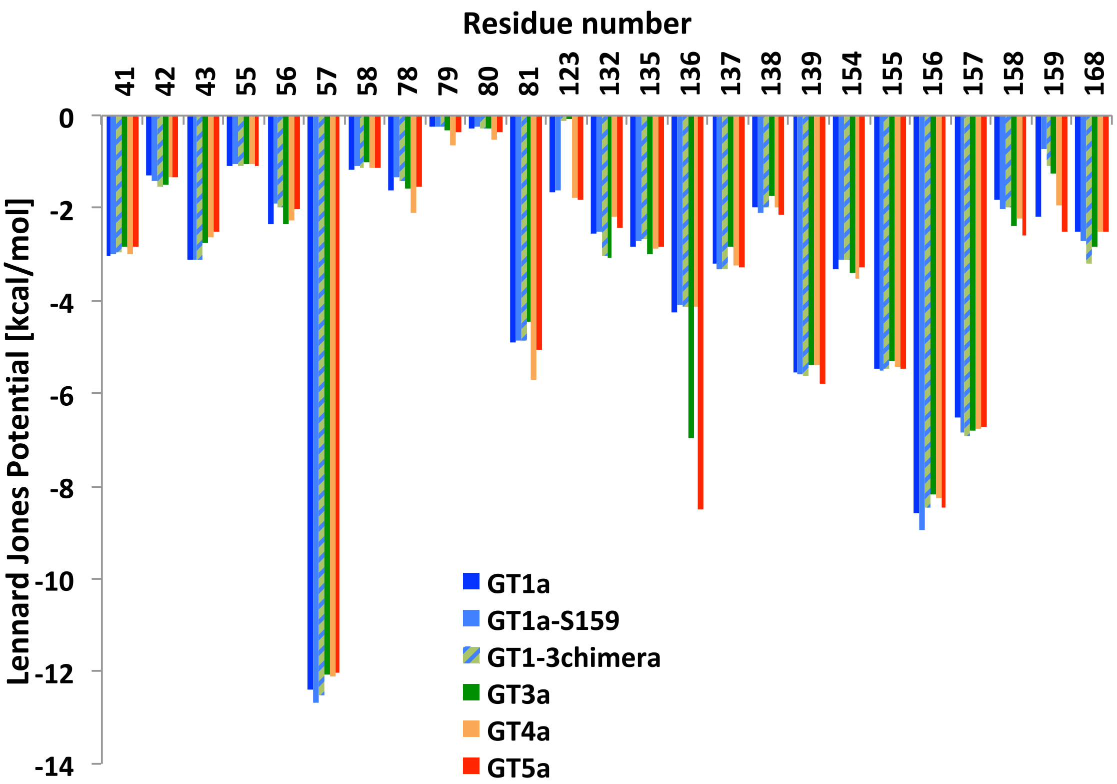


**Figure S.7: Van-der-Waals interactions between NS3/4A and GLE.** The vdW interactions between the different NS3/4A genotypes and GLE were calculated per residue and values above 0.5 kcal/mol were plotted.

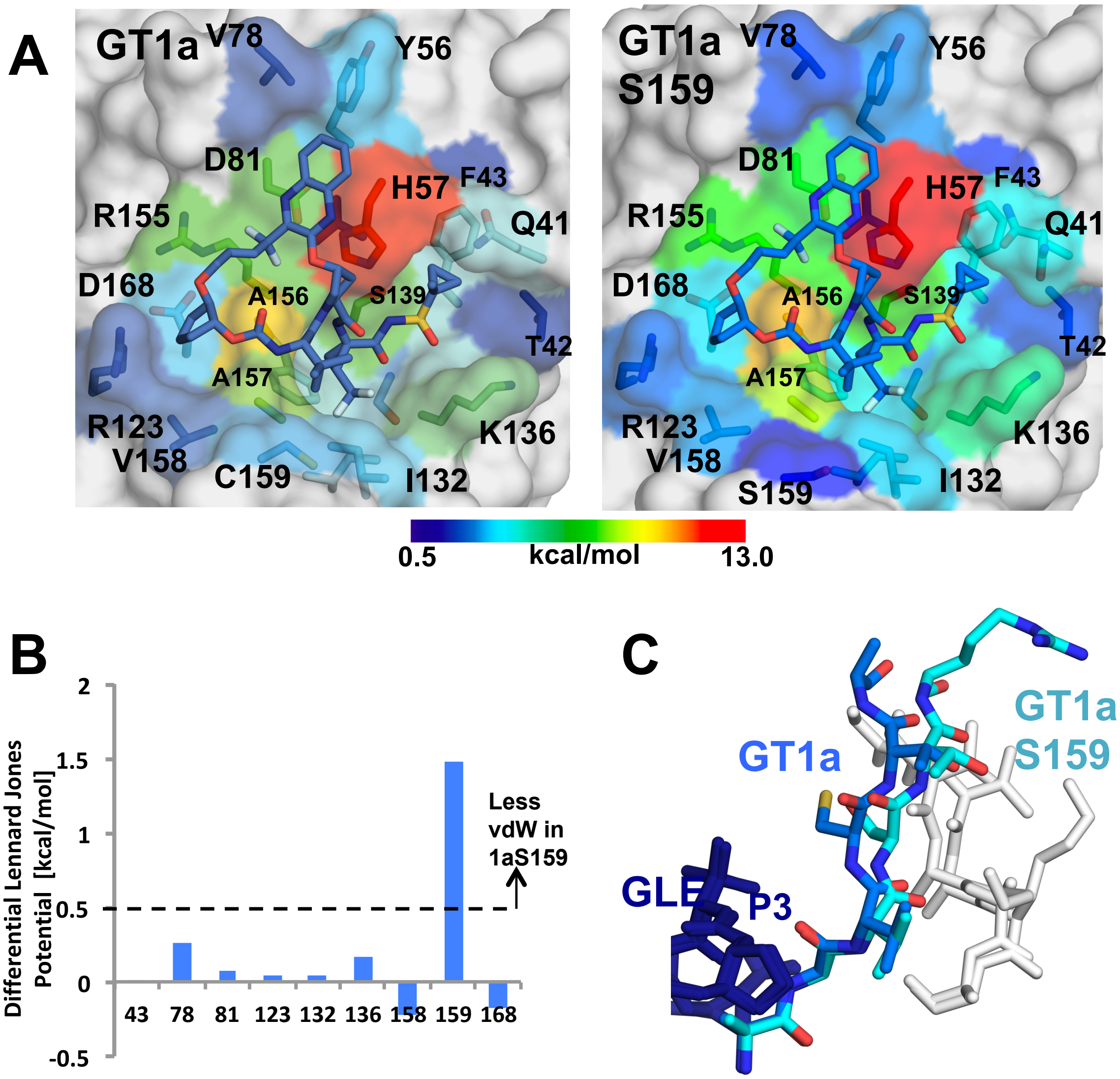


**Figure S.8: Van-der-Waals interactions between GT1a and GT1a-S159 and GLE.** (A) Structures of GT1a and GT1a-S159 in complex with GLE were aligned and displayed in Pymol. The Lennard Jones potentials (vdW) were displayed as color gradient on the calculated surface of GT1a and/or GT1a3a. All residues with vdW below 0.5 kcal/mol are shown in grey. (B) The differential vdW between the two structures were plotted as bar graph, only residue 159 showing changes in vdW above the threshold of 0.5 kcal/mol. (C) Overlay of GT1a and GT1a-S159 showing the conformational shift of the loop between residues 158 and163 responsible for the change in vdW interactions with GLE.
