## Supplementary material for "Molecular and structural mechanism of pan-genotypic HCV NS3/4A protease inhibition by glecaprevir": Table S.3

| | **GT1a_S159_GLE** | **GT1a_C159_GLE** | **GT1a_C159_D168E_GLE** | **GT1a3a_GRZ** | **GT1a3a_GLE** | **GT3a_GLE** | **GT4a_GLE** | **GT4a-SGGSGD_GLE** | **GT5a_GLE** |
|---|---|---|---|---|---|---|---|---|---|
| X-ray source | in house | in house | in house | APS beamline 23 ID-D | in house | APS beamline 23 ID-D | in house | in house | in house |
| Wavelength | 1.5418 | 1.5418 | 1.5418 | 1.0332 | 1.5418 | 1.0332 | 1.5418 | 1.5418 | 1.5418 |
| Resolution range | 26.359 - 1.728 (1.79 - 1.728) | 26.82 - 2.201 (2.28 - 2.201) | 20.38 - 2.002 (2.074 - 2.002) | 48.95 - 3.5 (3.625 - 3.5) | 26.341 - 1.749 (1.812 - 1.749) | 36.15 - 2.0 (2.072 - 2.0) | 30.49 - 2.3 (2.382 - 2.3) | 27.42 - 2.294 (2.376 - 2.294) | 21.33 - 2.0 (2.072 - 2.0) |
| Space group | P 21 21 21 | P 21 21 21 | P 21 21 21 | P 1 21 1 | P 21 21 21 | P 41 21 2 | P 21 21 21 | P 21 21 21 | P 21 21 21 |
| Unit cell (a, b, c, α, β, γ) | 55.026, 58.757, 60.058, 90, 90, 90 | 53.226, 58.55, 60.356, 90, 90, 90 | 55.157, 58.838, 60.488, 90, 90, 90 | 49.75, 58.968, 65.891, 90, 100.265, 90 | 54.571, 58.623, 60.053, 90, 90, 90 | 36.487, 36.487, 267.772, 90, 90, 90 | 44.093, 62.06, 60.989, 90, 90, 90 | 43.908, 60.738, 61.461, 90, 90, 90 | 39.551, 60.147, 83.788, 90, 90, 90 |
| Total reflections | 116126 (5283) | 57536 (3451) | 87043 (5613) | 9690 (949) | 194274 (4199) | 150951 (10392) | 50137 (2352) | 21565 (1384) | 98064 (9538) |
| Unique reflections | 19501 (1601) | 9722 (842) | 13619 (1293) | 4849 (475) | 19902 (1826) | 13303 (1255) | 7810 (735) | 7364 (659) | 13535 (1289) |
| Multiplicity | 5.9 (3.3) | 5.9 (4.1) | 6.4 (4.3) | 2.0 (2.0) | 9.7 (2.3) | 11.3 (8.3) | 6.4 (3.2) | 2.9 (2.1) | 7.2 (7.4) |
| Completeness (%) | 92.56 (77.27) | 97.04 (87.33) | 98.80 (95.42) | 99.86 (99.58) | 99.23 (93.45) | 99.91 (99.92) | 99.46 (96.07) | 94.71 (86.37) | 96.05 (93.55) |
| Mean I/sigma(I) | 37.76 (5.7) | 13.24 (3.38) | 9.33 (2.31) | 7.18 (4.50) | 30.94 (4.09) | 13.98 (2.25) | 22.01 (3.23) | 15.44 (3.4) | 44.17 (14.83) |
| Wilson B-factor | 14.6 | 23.06 | 20.39 | 32.92 | 15.69 | 32.64 | 26.6 | 25.97 | 19.27 |
| R-merge | 0.045 (0.180) | 0.1066 (0.3488) | 0.1649 (0.5775) | 0.08193 (0.1376) | 0.067 (0.165) | 0.1236 (0.7479) | 0.080 (0.302) | 0.065 (0.217) | 0.045 (0.140) |
| R-meas | | 0.1168 (0.3969) | 0.1793 (0.6527) | 0.1159 (0.1946) | | 0.1295 (0.7982) | | | |
| R-pim | | 0.04683 (0.1844) | 0.06947 (0.296) | 0.08193 (0.1376) | | 0.03796 (0.2718) | | | |
| CC1/2 | | 0.996 (0.919) | 0.995 (0.785) | 0.984 (0.963) | | 0.997 (0.563) | | | |
| CC* | | 0.999 (0.979) | 0.999 (0.938) | 0.996 (0.99) | | 0.999 (0.849) | | | |
| Reflections used in refinement | 19442 (1601) | 9718 (841) | 13614 (1293) | 4847 (474) | 19895 (1826) | 13293 (1254) | 7803 (734) | 7364 (659) | 13511 (1290) |
| Reflections used for R-free | 975 (67) | 485 (46) | 678 (67) | 243 (24) | 2010 (183) | 665 (62) | 781 (76) | 366 (33) | 677 (65) |
| R-work | 0.1526 (0.1779) | 0.2058 (0.2423) | 0.1718 (0.2165) | 0.2205 (0.2329) | 0.1397 (0.1772) | 0.2087 (0.2968) | 0.1697 (0.1906) | 0.1925 (0.2468) | 0.1656 (0.1791) |
| R-free | 0.1845 (0.2162) | 0.2558 (0.2984) | 0.2169 (0.2239) | 0.2626 (0.2635) | 0.1797 (0.2112) | 0.2365 (0.3190) | 0.2166 (0.2252) | 0.2422 (0.2858) | 0.2127 (0.2148) |
| CC(work) | | 0.949 (0.888) | 0.966 (0.869) | 0.900 (0.913) | | 0.868 (0.506) | | | |
| CC(free) | | 0.910 (0.804) | 0.945 (0.943) | 0.835 (0.918) | | 0.754 (0.650) | | | |
| Number of non-hydrogen atoms | 1924 | 1663 | 1827 | 2945 | 1882 | 1571 | 1367 | 1271 | 1772 |
| macromolecules | 1552 | 1478 | 1550 | 2723 | 1508 | 1408 | 1168 | 1134 | 1481 |
| ligands | 85 | 71 | 71 | 169 | 78 | 71 | 59 | 59 | 64 |
| solvent | 287 | 114 | 206 | 53 | 296 | 92 | 140 | 78 | 227 |
| Protein residues | 200 | 202 | 203 | 393 | 203 | 192 | 159 | 160 | 193 |
| RMS(bonds) | 0.009 | 0.003 | 0.002 | 0.002 | 0.008 | 0.005 | 0.003 | 0.002 | 0.006 |
| RMS(angles) | 0.9 | 0.56 | 0.58 | 0.61 | 0.91 | 0.7 | 0.57 | 0.53 | 0.64 |
| Ramachandran favored (%) | 98.48 | 97.99 | 98.48 | 94.06 | 98.99 | 97.37 | 96.77 | 96.75 | 97.38 |
| Ramachandran allowed (%) | 1.52 | 2.01 | 1.52 | 5.68 | 1.01 | 2.63 | 3.23 | 3.25 | 2.62 |
| Ramachandran outliers (%) | 0 | 0 | 0 | 0.26 | 0 | 0 | 0 | 0 | 0 |
| Rotamer outliers (%) | 2.38 | 0.64 | 0 | 0 | 0 | 4.2 | 1.57 | 2.46 | 1.88 |
| Clashscore | 5.27 | 5.04 | 3.16 | 4.91 | 6.79 | 5.95 | 2.89 | 3 | 3.63 |
| Average B-factor | 19.73 | 25.84 | 24.11 | 32.38 | 21.06 | 50.21 | 29.63 | 26.32 | 23.59 |
| macromolecules | 17.54 | 25.48 | 23.1 | 32.05 | 18.44 | 50.97 | 29.2 | 26.47 | 22.6 |
| ligands | 20.74 | 24.56 | 24.73 | 42.12 | 19.79 | 36.03 | 21.64 | 20.87 | 17.96 |
| solvent | 31.28 | 31.19 | 31.42 | 18.27 | 34.76 | 49.46 | 36.56 | 28.31 | 31.64 |
| Molprobity score | 1.57 | 1.34 | 1.11 | 1.66 | 1.37 | 1.89 | 1.41 | 1.58 | 1.43 |
| **PDB ID** | **6P6L** | **6P6M** | **6P6O** | **6P6Q** | **6P6R** | **6P6S** | **6P6T** | **6P6Z** | **6P6V** |
